# SARS-CoV-2 diverges from other betacoronaviruses in only partially activating the IRE1α/XBP1 ER stress pathway in human lung-derived cells

**DOI:** 10.1101/2021.12.30.474519

**Authors:** Long C. Nguyen, David M. Renner, Diane Silva, Dongbo Yang, Nicholas Parenti, Kaeri M. Medina, Vlad Nicolaescu, Haley Gula, Nir Drayman, Andrea Valdespino, Adil Mohamed, Christopher Dann, Kristin Wannemo, Lydia Robinson-Mailman, Alan Gonzalez, Letícia Stock, Mengrui Cao, Zeyu Qiao, Raymond E. Moellering, Savas Tay, Glenn Randall, Michael F. Beers, Marsha Rich Rosner, Scott A. Oakes, Susan R. Weiss

## Abstract

Severe acute respiratory syndrome coronavirus 2 (SARS-CoV-2) has killed over 6 million individuals worldwide and continues to spread in countries where vaccines are not yet widely available, or its citizens are hesitant to become vaccinated. Therefore, it is critical to unravel the molecular mechanisms that allow SARS-CoV-2 and other coronaviruses to infect and overtake the host machinery of human cells. Coronavirus replication triggers endoplasmic reticulum (ER) stress and activation of the unfolded protein response (UPR), a key host cell pathway widely believed essential for viral replication. We examined the master UPR sensor IRE1α kinase/RNase and its downstream transcription factor effector XBP1s, which is processed through an IRE1α-mediated mRNA splicing event, in human lung-derived cells infected with betacoronaviruses. We found human respiratory coronavirus OC43 (HCoV-OC43), Middle East respiratory syndrome coronavirus (MERS-CoV), and murine coronavirus (MHV) all induce ER stress and strongly trigger the kinase and RNase activities of IRE1α as well as XBP1 splicing. In contrast, SARS-CoV-2 only partially activates IRE1α through autophosphorylation, but its RNase activity fails to splice XBP1. Moreover, while IRE1α was dispensable for replication in human cells for all coronaviruses tested, it was required for maximal expression of genes associated with several key cellular functions, including the interferon signaling pathway, during SARS-CoV-2 infection. Our data suggest that SARS-CoV-2 actively inhibits the RNase of autophosphorylated IRE1α, perhaps as a strategy to eliminate detection by the host immune system.

**IMPORTANCE:** SARS-CoV-2 is the third lethal respiratory coronavirus after MERS-CoV and SARS-CoV to emerge this century, causing millions of deaths world-wide. Other common coronaviruses such as HCoV-OC43 cause less severe respiratory disease. Thus, it is imperative to understand the similarities and differences among these viruses in how each interacts with host cells. We focused here on the inositol-requiring enzyme 1α (IRE1α) pathway, part of the host unfolded protein response to virus-induced stress. We found that while MERS-CoV and HCoV-OC43 fully activate the IRE1α kinase and RNase activities, SARS-CoV-2 only partially activates IRE1α, promoting its kinase activity but not RNase activity. Based on IRE1α-dependent gene expression changes during infection, we propose that SARS-CoV-2 prevents IRE1α RNase activation as a strategy to limit detection by the host immune system.

## INTRODUCTION

Severe acute respiratory syndrome coronavirus 2 (SARS-CoV-2) emerged in China in late 2019. It was the third lethal zoonotic coronavirus to emerge into humans after SARS-CoV (2002) and Middle East respiratory syndrome coronavirus (MERS-CoV) (2012), each of which has been associated with acute lung injury and hypoxemic respiratory failure. While coronaviruses are divided into four genera (alpha, beta, gamma, and delta)(1, 2), all three of the lethal human coronaviruses are betacoronaviruses, albeit from different lineages (Figure 1). SARS-CoV and SARS-CoV-2 are sarbecoviruses, while MERS-CoV is a merbecovirus. Other human CoVs, including HCoV-OC43 (OC43) and HCoV-HKU1 (HKU-1), are embecoviruses as is the model murine coronavirus mouse hepatitis virus (MHV). All CoVs have similar genome structures, replication cycles, and the human CoVs as well as some MHV strains exhibit tropism for the epithelia of the respiratory tract, the portal of entry. They replicate their RNAs and produce subgenomic mRNAs by conserved mechanisms and encode homologous structural as well as replicase proteins. Despite the similarities among all coronaviruses, each lineage expresses distinct accessory proteins that may confer differences in host-virus interactions. Indeed, we have previously found that SARS-CoV-2, MERS-CoV and MHV all induce somewhat different levels of activation and/or antagonism of interferon (IFN) signaling and other dsRNA induced antiviral innate responses (3-5).

**Figure 1.**
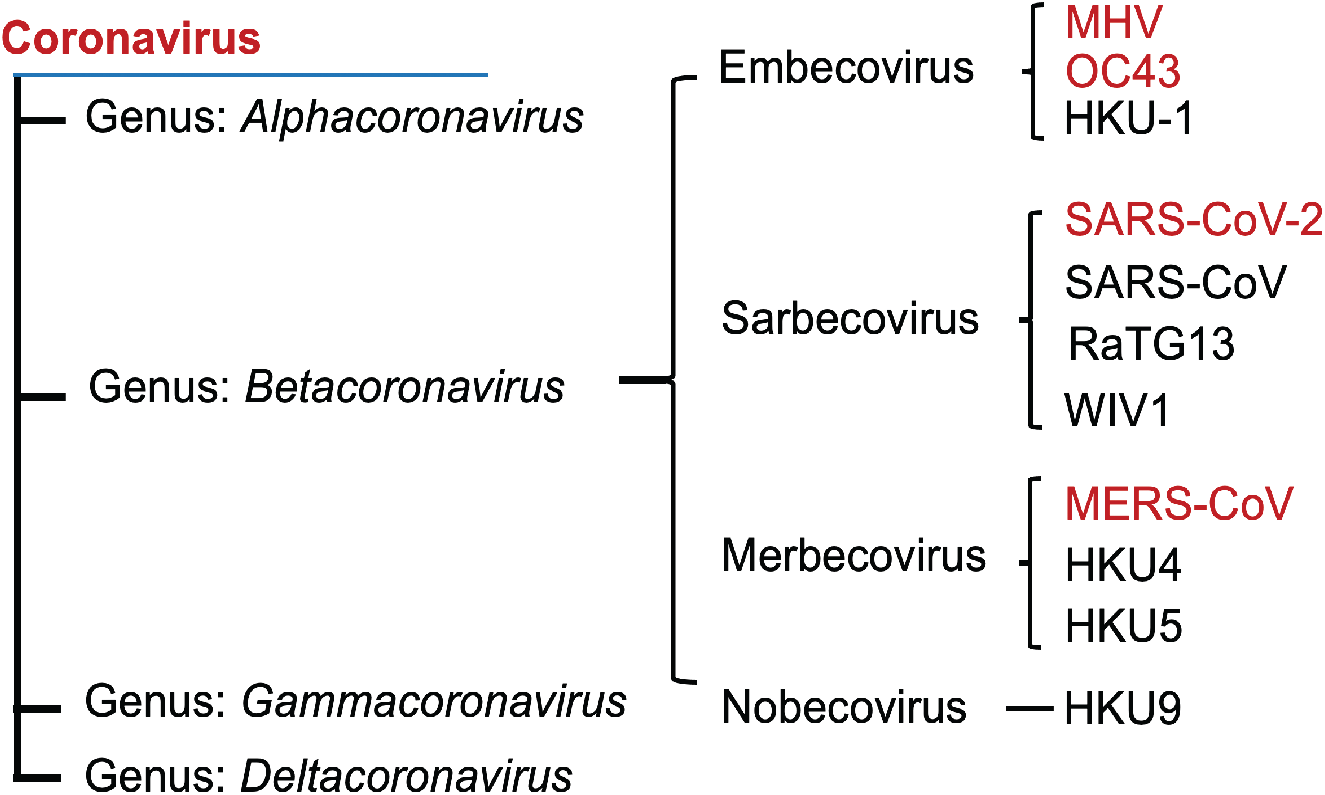
Coronavirus family. Phylogenetic tree of betacoronaviruses and their lineages. Viruses examined in this study are show in red font.

One key pathway involved in the virus-induced host response is the endoplasmic reticulum (ER) stress response that regulates protein homeostasis (referred to as proteostasis) in this organelle. One third of all eukaryotic proteins, including most that are inserted into membranes or secreted, are synthesized through co-translational translocation into the ER lumen. Likewise, viral membrane associated proteins are translated and processed in association with the ER (6, 7). Once in the ER, these polypeptides undergo stringent quality control monitoring to ensure that they are properly processed and folded. If the capacity to fold proteins is unable to keep up with demand, misfolded proteins will accumulate in the ER lumen—a condition referred to as “ER stress.” The presence of misfolded proteins in the ER is sensed by three transmembrane sentinel proteins - activating transcription factor 6 (ATF6), PKR-like ER kinase (PERK), and inositol-requiring enzyme (IRE)1α - which trigger an intracellular signaling pathway called the unfolded protein response (UPR). In an effort to restore proteostasis, activation of these sensors induces transcription factors that turn on genes encoding chaperones, oxidoreductases, and ER-associated decay (ERAD) components(8). The UPR also inhibits cap-dependent translation, thus decreasing the load on the ER and giving it extra time to fold proteins already in production (9, 10). If successful, these adaptive UPR programs restore ER homeostasis.

The most ancient UPR pathway is controlled by IRE1α — an ER transmembrane bifunctional kinase/endoribonuclease (RNase) that employs auto-phosphorylation to control its catalytic RNase function (11, 12). In response to ER stress, IRE1α undergoes auto-phosphorylation and dimerization to allosterically activate its RNase domain to excise a 26nt non-conventional intron in *XBP1* mRNA; re-ligation of spliced *XBP1* shifts the open reading frame, and its translation produces the homeostatic transcription factor XBP1s (s=spliced) (13, 14). Once synthesized, XBP1s upregulates genes that expand the ER and its protein folding machinery (15). IRE1α can additionally lead to apoptosis and inflammation via JUN N-terminal kinase (JNK) and p38 mitogen-activated protein kinase (MAPK) signaling (16). Prolonged ER stress can induce regulated IRE1-dependent decay (RIDD), promoting the cleavage of additional targets beyond XBP1 mRNA, such as secretory protein and ER-localized mRNAs (17). In the short term, RIDD may promote adaptation through further reducing translation and protein burden on the ER. However, prolonged RIDD leads to the depletion of vital ER resident enzymes and structural components to exacerbate ER stress and hasten cell death (11, 18).

There is a large body of evidence that viral replication of mammalian cells can trigger ER stress and UPR activation in infected cells (19), and numerous studies report that the UPR is activated upon infection of host cells by coronavirus family members (6, 7, 20-25) Coronaviruses induce stress in the ER in several ways. First, conserved replicase encoded, nonstructural proteins nsp3, nsp4 and nps6 are embedded into the ER membrane, and along with unknown host factors, promote membrane curvature to form double membrane vesicles (DMVs), the site of viral replication/transcription centers (RTC) (26). In addition to remodeling the ER, coronaviruses further condition infected cells by shifting translation away from host mRNAs and instead to viral mRNAs. Translation of viral mRNAs causes the ER to be flooded with heavily glycosylated viral structural proteins [e.g., spike (S), membrane (M) and envelope (E)], challenging the organelle’s folding capacity and overall integrity. Indeed, overexpression of CoV spike proteins (27) as well as several sarbecovirus accessory poteins (22, 28) has been reported to induce ER stress. Finally, cell membranes are depleted as enveloped virus particles are assembled into new virions in the ER-Golgi intermediate compartment before budding from the infected cell (1). Thus, coronaviruses as well as other enveloped viruses promote a massive ER expansion and modification necessary to replicate their genomes, transcribe mRNAs, and finally to process and package their protein products into viral particles.

We have compared the activation status and requirement of the IRE1α/XBP1 arm of the UPR in well-characterized human lung epithelial cell lines and in induced pluripotent stem cell (iPSC)-derived type II alveolar (iAT2) cells, following infection with four betacoronaviruses representing three distinct lineages. We find that infection with MERS-CoV, OC43 and MHV leads to phosphorylation of IRE1α and the consequent production of spliced XBP1 transcription factor. Surprisingly, while we observed phosphorylation of IRE1α in SARS-CoV-2 infected cells, there was notable absence of XBP1s, suggesting SARS-CoV-2 inhibits downstream signaling of the IRE1α/XBP1 arm of the UPR. In addition, we report reduced SARS-CoV-2 induced interferon signaling gene expression in the absence of IRE1α.

## RESULTS

### Induction of IRE1α phosphorylation following coronavirus infection

To determine whether betacoronaviruses activate IRE1α, we first examined the level of phosphorylated IRE1α after viral infection of the A549 human lung carcinoma cell line. We used A549 cells stably expressing the following receptors to facilitate optimal entry for each of the viruses: carcinoembryonic antigen cell adhesion molecule (CEACAM)1a or MHVR (MHV), dipeptidyl peptidase DPP4 (MERS-CoV), or angiotensin converting enzyme (ACE)2 (SARS-CoV-2). HCoV-OC43 can infect parental A549 or cells expressing ACE2 (3). Consistent with previous reports that embeco lineage coronaviruses MHV (20, 29) and OC43 (24) induce ER stress, we observed a significant increase in phospho-IRE1α (p-IRE1α) during infection by either OC43 (24 or 48hpi) or MHV (24hpi) (Figure 2A-C). To confirm the specificity of the p-IRE1α band, we pretreated cells prior to infection with KIRA8, a highly selective kinase inhibitor of IRE1α known to inhibit both autophosphorylation and consequently RNase activity. As expected, KIRA8 significantly inhibited the induction of p-IRE1α by OC43 and MHV (Figure 2A&C). Thapsigargin (Tg) and tunicamycin (Tm), both inducers of ER stress, were used as further controls (Figure 2B,D&E). Robust induction of p-IRE1α was observed with 1 hour of Tg (1μM) treatment, while no activation of p-IRE1α was observed after 8 hours of treatment with Tm (1μg/ mL), consistent with the negative feedback regulation observed with extended Tm treatment (30). We also observed robust phosphorylation of IRE1α in A549-DDP4 cells and A549-ACE2 cells infected by MERS-CoV and SARS-CoV-2, respectively at 24 and 48 hpi (Figures 2D-F and S1A&B). As with OC43 and MHV, IRE1α phosphorylation during SARS-CoV-2 infection was inhibited by KIRA8 (Figure 2F). These results are not limited to a single cell type as we observed similar induction of p-IRE1α in Calu-3 cells, another lung epithelial derived cells line, which can be productively infected with both MERS-CoV or SARS-CoV-2 (Figure 2G). These results demonstrate that MERS-CoV, SARS-CoV-2, HCoV-OC43 and MHV activate the host IRE1α kinase after infection.

**Figure 2.**
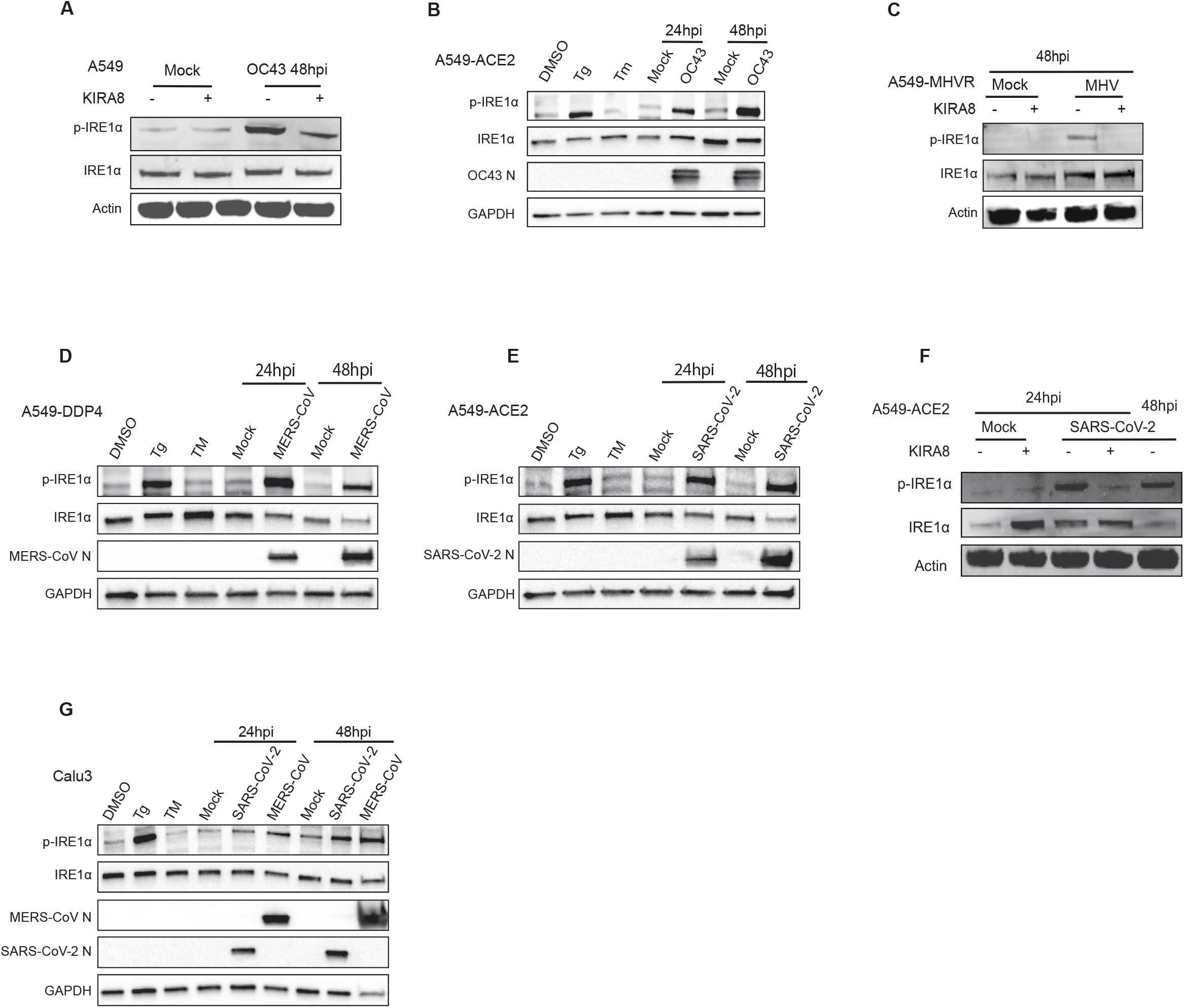
Induction of IRE1α phosphorylation following coronavirus infection. A549 cells expressing the indicated viral receptors were mock infected or infected. Protein was harvested at 24 or 48hpi and analyzed by immunoblotting with antibodies, as indicated. (A,C,F) Cells infected with OC43 at MOI=4 (A) or MHV at MOI=0.1 (C) or SARS-CoV-2 at MOI=3 (F) were pre-treated 2 hours prior to infection with 1μM KIRA8. (B,D,E) Cells were infected with OC43 at MOI=1 (B), MERS-CoV at MOI=5 (D), or SARS-CoV-2 at MOI=5 (E) or treated with DMSO, thapsigargin (Tg, 1μM) for 1 hour or tunicamycin (TM, 1μg/ mL) for 8 hours. (G) Calu-3 cells were mock infected, or infected with MERS-CoV, or SARS-CoV-2 (MOI=5).. Data shown are from one representative of at least two independent experiments.

### MHV, OC43, MERS-CoV but not SARS-CoV-2 induce splicing of XBP1 mRNA

We next examined the effect of coronavirus infection on the RNase activity of IRE1α as assessed by XBP1 splicing. Using specific primers to quantify spliced XBP1 mRNA (XBP1s), we observed a marked increase in the percentage of spliced XBP1 mRNA (% XBP1s) as well as an increase in the relative amount of spliced XBP1 mRNA (XBP1s) compared to mock control after infection by OC43, MERS-CoV or MHV in receptor-expressing A549 cells (Figures 3A&B and S2A&B). This induction of XBP1s by OC43 and by MERS-CoV infection was confirmed by assessing XBP1 splicing by agarose gel electrophoresis (Figure 3E&F). DNAJB9, a canonical target of XBP1s, was also markedly upregulated with OC43, MERS-CoV, and MHV infection at both 24 and 48 hours post-infection (Figures 3A&B and S2B). This induction of IRE1α RNase activity is coincident with the observed autophosphorylation of p-IRE1α upon OC43, MHV or MERS-CoV infection.

**Figure 3.**
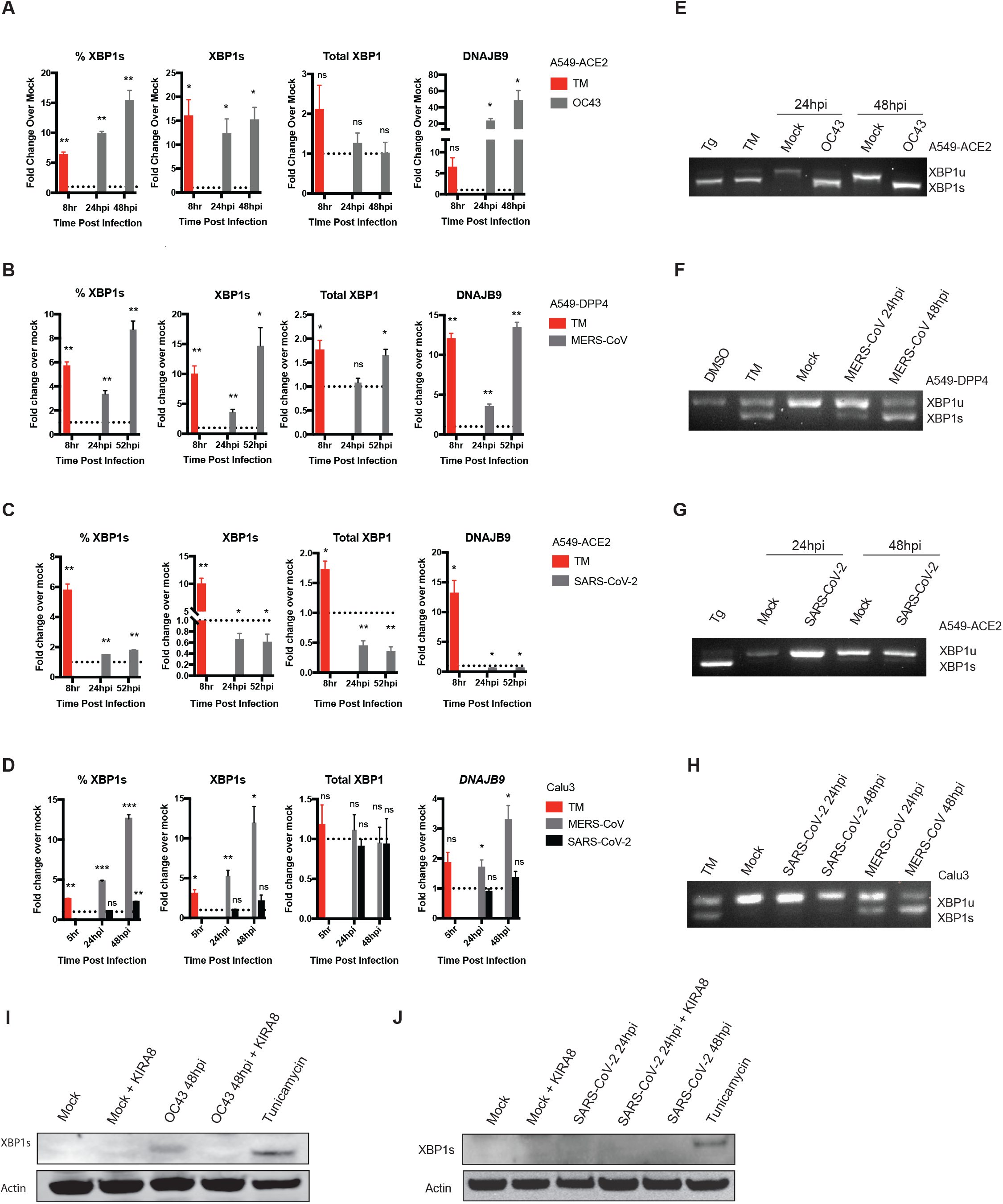
IRE1α-mediated XBP1 splicing occurs following infection with OC43 or MERS-CoV, but not SARS-CoV-2. A549 cells were mock infected or infected (in triplicate) with OC43 at MOI=1 (A, E), MERS-CoV at MOI=5 (B, F), SARS-CoV-2 at MOI=5 (C, G) or treated with Tm (1μg/ mL) for 8 hours and total RNA harvested at indicated time points. (A-C) Relative %XBP1s, XBP1s, total XBP1 and DNAJB9 mRNA expression were quantified by RT-qPCR. C_T_ values were normalized to 18S rRNA and expressed as fold-change over mock displayed as 2^−Δ(ΔCt)^. Technical replicates were averaged, the means for each replicate displayed, ±SD (error bars). (D) Calu-3 cells were mock infected or infected with MERS-CoV or SARS-CoV-2 (MOI=5) and total RNA harvested at indicated time points. Relative %XBP1s, XBP1s, total XBP1 and DNAJB9 mRNA expression were quantified by RT-qPCR, calculated, and displayed as described above. Values are means ± SD (error bars). Statistical significance was determined using two-tailed, paired Student’s *t*-test. Displayed significance (infected relative to mock) is determined by p-value (P), where * = P < 0.05; ** = P < 0.01; *** = P < 0.001; **** = P < 0.0001; ns = not significant. (E-H) RNA was harvested from A549 cells mock infected or infected with OC43 at MOI=1 (E), MERS-CoV at MOI=5 (F), SARS-CoV-2 at MOI=5 (G), or Calu-3 cells infected with MERS-CoV and SARS-CoV-2 at MOI=5 (H) or treated with tunicamycin (Tm, 1μg/ mL) for 8 hour, or thapsigargin (Tg, 1μM) for 1 hour or DMSO. RT-PCR was performed using primers crossing the XBP1 splicing site. The product was resolved on an agarose gel to visualize XBP1 splicing. (I-J) Lysates from A549-ACE2 cells mock infected, or Tm (500 ng/mL) for 6 hours or infected with OC43 (MOI=4) or SARS-CoV-2 (MOI=3), treated with or without KIRA8 (1μM), were harvested at indicated time points as in Figure 2A,C&F and immunoblotted with antibody directed against XBP1s protein. Data shown are from one representative experiment from at least three independent experiments.

Surprisingly, despite the observed IRE1α autophosphorylation following SARS-CoV-2 infection, there was no significant upregulation of XBP1s mRNA in A549-ACE2 cells up to 52 hours post-infection (Figure 3C&G). Similarly, DNAJB9 expression levels were unchanged at all time points observed with SARS-CoV-2 (Figure 3C). To confirm this effect is not limited to A549 cells, we measured XBP1 mRNA splicing in MERS-CoV and SARS-CoV-2-infected Calu-3 cells. Again, infection with MERS-CoV, but not SARS-CoV-2, significantly induced XBP1s and its downstream effector DNAJB9 (Figure 3D&H). In agreement with these results, OC43, but not SARS-CoV-2, infection induced XBP1s protein levels (Figure 3I&J).

### Upon infection, MHV, OC43, MERS-CoV induce IRE1α and related genes to a greater extent than SARS-CoV-2

To determine how different coronaviruses impact the UPR at the transcriptional level, we performed RNA-sequencing of A549-DPP4 cells infected with MERS-CoV for 24 and 36 hours. We compared the results to published RNA-seq data sets (29, 31) of MHV infection of murine bone marrow derived macrophages (BMDM) or SARS-CoV-2 infection of A549-ACE2, normal human bronchial epithelial (NHBE) cells, and Calu-3 cell lines. In agreement with our IRE1α activation results, Ingenuity Pathway Analysis (IPA) predicted activation of the UPR and ER stress pathways by MERS-CoV and MHV (Figure 4A). In contrast, SARS-CoV-2 consistently showed little to no activation of the UPR and ER stress pathway across different MOI conditions and cell lines.

**Figure 4.**
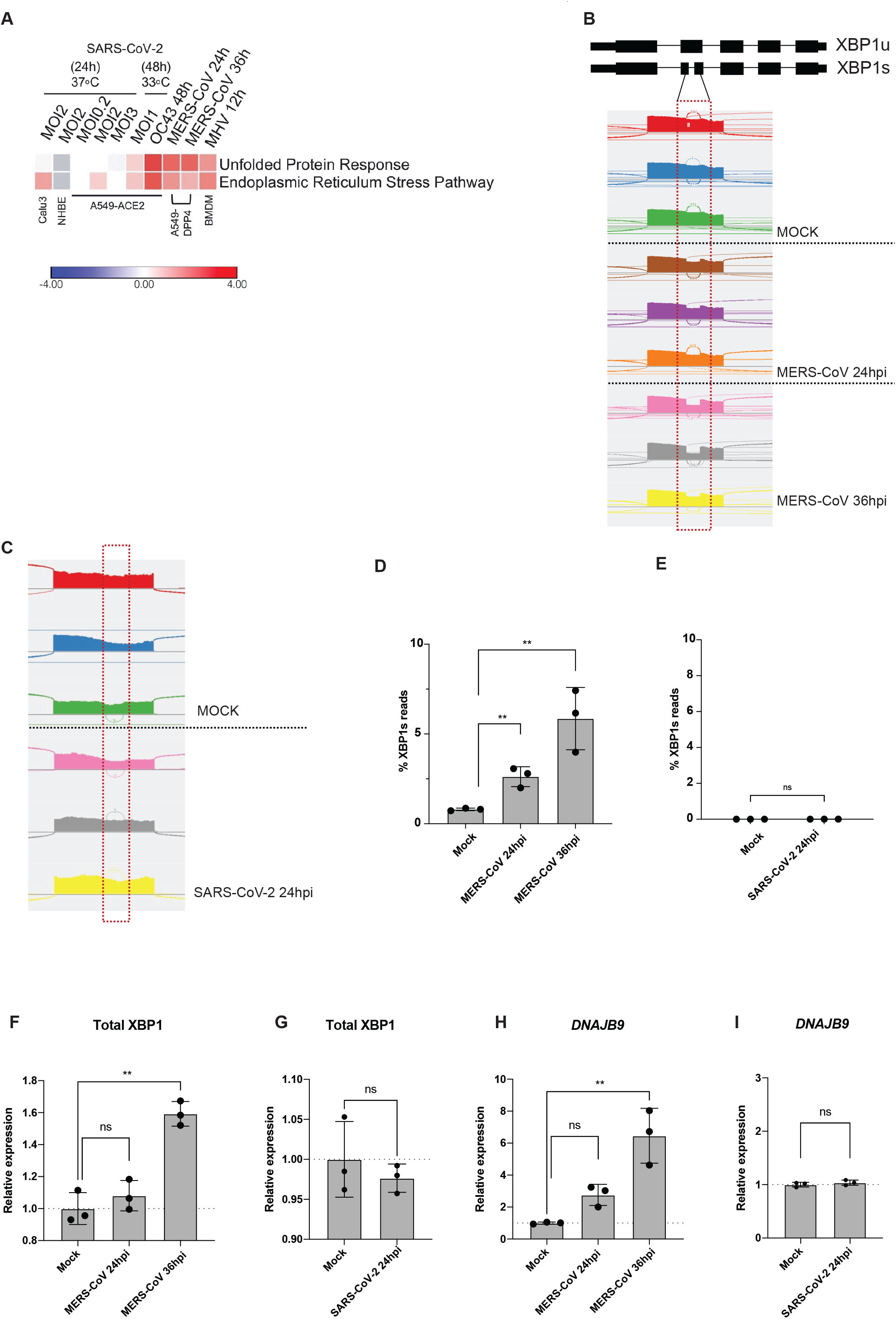
Unlike other coronaviruses, SARS-CoV-2 infection does not lead to robust UPR activation. (A) Heatmap of predicted pathway status based on Ingenuity Pathway Analysis (IPA) of activation z-scores for each pathway from RNA-sequencing data from indicated cells infected with OC43 (MOI =1), MERS-CoV (MOI = 1), MHV (MOI = 1) and SARS-CoV-2 under specified conditions. Red: pathway predicted to be activated. Blue: pathway predicted to be inhibited. White: pathway predicted to be unchanged. Gray: no prediction due to lack of significance. (B&C) Quantification of XBP1 splicing by analyzing RNA-Seq data from A549-DPP4 and A549-ACE2 cells mock-infected or infected with MERS-CoV or SARS-CoV-2, respectively, under indicated conditions. Reads representing spliced or unspliced XBP1 mRNA were identified based on the presence or absence of the 26 nucleotides intron and quantified. (D-I) Percentage of XBP1 spliced reads, or relative expression of total XBP1 and DNAJB9 mRNA from the RNA-seq samples. Values are means ± SD (error bars). Statistical significance was determined by Unpaired t-tests (* = P < 0.05; ** = P < 0.01; ns = not significant).

To support the results of the gel electrophoresis splicing assays for XBP1 mRNA that distinguished SARS-CoV-2 infection from that of the other betacoronaviruses (Figure 3), we further utilized the RNA sequencing results to quantitatively measure XBP1 mRNA splicing by these coronaviruses. Through RNA-seq, we visualized both the unspliced and spliced XBP1 mRNA reads based on whether they contain the 26 nucleotide non-conventional intron that is removed as a result of RNase activity of IRE1α as previously described (32) (Figure 4B&C). MERS-CoV infection resulted in significant XBP1 mRNA splicing, in contrast with no difference detected in SARS-CoV-2 infected versus mock-infected cells (Figure 4B&C). We further quantified total XBP1 spliced vs unspliced reads, which consistently showed a substantial increase in the percent expression of the XBP1s reads when normalized to total XBP1 reads for MERS-CoV at both 24 and 36 hours post-infection but not for SARS-CoV-2 infected cells (Figure 4D&E). This was consistent with significant upregulation of DNAJB9 and total XBP1 during infection with MERS-CoV but not SARS-CoV-2 (Figure 4F-I).

### MERS-CoV but not SARS-CoV-2 induces XBP1 splicing during infection of biologically relevant iPSC-derived alveolar type II cells

To confirm our results in a more physiologically relevant cell, we infected iPSC-derived type II alveolar (iAT2) cells. We employed the SPC2 line, which expresses tdTomato from the surfactant protein-C (SFTPC) locus as an AT2 marker, which we have previously used to characterize innate immune responses to SARS-CoV-2 infection (3). Type II alveolar cells are a major target during both MERS-CoV and SARS-CoV-2 infection in humans, and their destruction may be a contributing factor to lung pathogenesis in severe cases (33, 34).

Both MERS-CoV and SARS-CoV-2 replicate in these cells and release infectious virus as quantified by plaque assay (Figure 5A). Notably, MERS-CoV replicated to higher titers than SARS-CoV-2 in these lung-derived cells. This complements our previous findings that SARS-CoV-2 replicates more efficiently than MERS-CoV in upper respiratory derived primary nasal cells (3), and may suggest that MERS-CoV is better adapted to replicate within the lower respiratory tract while SARS-CoV-2 replicates more efficiently in the upper airway. Despite this difference in replication, both viruses were observed to induce p-IRE1α over the course of infection (Figure 5B). In agreement with our results in A549 and Calu-3 cells, SARS-CoV-2 failed to induce XBP1 splicing in iAT2 cells, as measured by RT-qPCR (Figure 5C). By contrast, MERS-CoV induced XBP1 splicing, albeit to a lower extent than in immortalized cell lines. Lastly, we visualized XBP1 splicing using RT-PCR and agarose gel electrophoresis (Figure 5D). Again, our data indicate that SARS-CoV-2 fails to induce XBP1 splicing at either 24 or 48hpi in iAT2 cells, despite inducing p-IRE1α. MERS-CoV, however, induced increasing XBP1 splicing over the course of infection, matching the results in A549 and Calu-3 cells (Figures 2 and 3). Overall, these results indicate that both SARS-CoV-2 and MERS-CoV induce ER stress as evidenced by IRE1α phosphorylation during infection of primary iAT2 cells, but only MERS-CoV induces the downstream effects of active IRE1α RNase.

**Figure 5.**
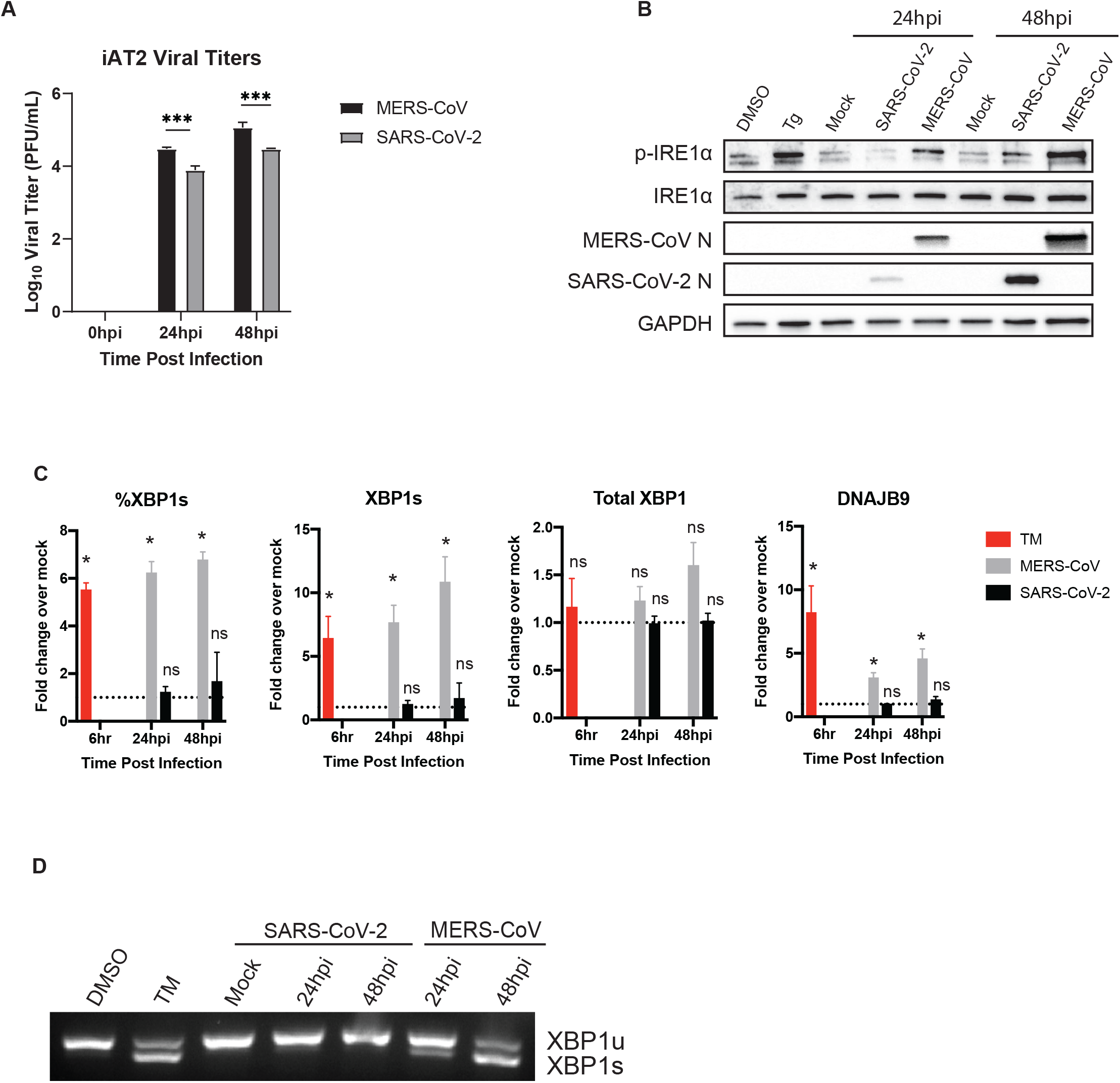
SARS-CoV-2 and MERS-CoV induce IRE1α phosphorylation in iAT2 cells but diverge in induction of XBP1 splicing. iPSC-derived AT2 cells (iAT2 cells) were mock infected or infected (in triplicate) with MERS-CoV or SARS-CoV-2 at a MOI of 5. (A) At the indicated timepoints, supernatants were collected, and infectious virus quantified by plaque assay. Values are means ± SD (error bars). Statistical significance was determined by two-way ANOVA (* = P < 0.05; ns = not significant). (B) Total protein was harvested at the indicated timepoints and analyzed by immunoblotting using the indicated antibodies. Thapsigargin treatment for 1 hour (Tg; 1μM) was used as a positive control for IRE1α activation while DMSO served as a vehicle control. (C) Total RNA was harvested at the indicated timepoints and relative %XBP1s, XBP1s, and total XBP1 mRNA expression were quantified by RT-qPCR, calculated, and displayed as described above. Values are means ± SD (error bars). Statistical significance (infected compared to mock) was determined using two-tailed, paired Student’s *t*-test. Displayed significance is determined by p-value (P), where * = P < 0.05; ** = P < 0.01; *** = P < 0.001; **** = P < 0.0001; ns = not significant. (D) RT-PCR was performed using extracted RNA and primers crossing the XBP1 splicing site. The product was run out on an agarose gel to visualize XBP1 splicing. Tunicamycin treatment (1μg/mL for 6 hours) was used as a positive control for RT-(q)PCR, while DMSO treatment served as a vehicle control. Data shown are from one representative experiment from at least two independent experiments.

### SARS-CoV-2 inhibits XBP1 splicing

We then tested whether SARS-CoV-2 actively inhibits splicing of XBP1 induced by the N-linked glycosylation inhibitor tunicamycin (TM), a common agent used to chemically induce ER stress. To do so, A549-ACE2 cells were either mock infected or infected with SARS-CoV-2 or OC43 for 24 hours and then treated with TM for 6 hours prior to analysis. Interestingly, while SARS-CoV-2 infection did not completely prevent XBP1 splicing induced by TM, it led to significantly lower XBP1 splicing levels compared with mock infected cells (Figure 6A). In contrast, OC43 increased XBP1 splicing at all tested concentrations of TM (Figure 6B). This result suggests that SARS-CoV-2 actively inhibits activation IRE1α RNase.

**Figure 6.**
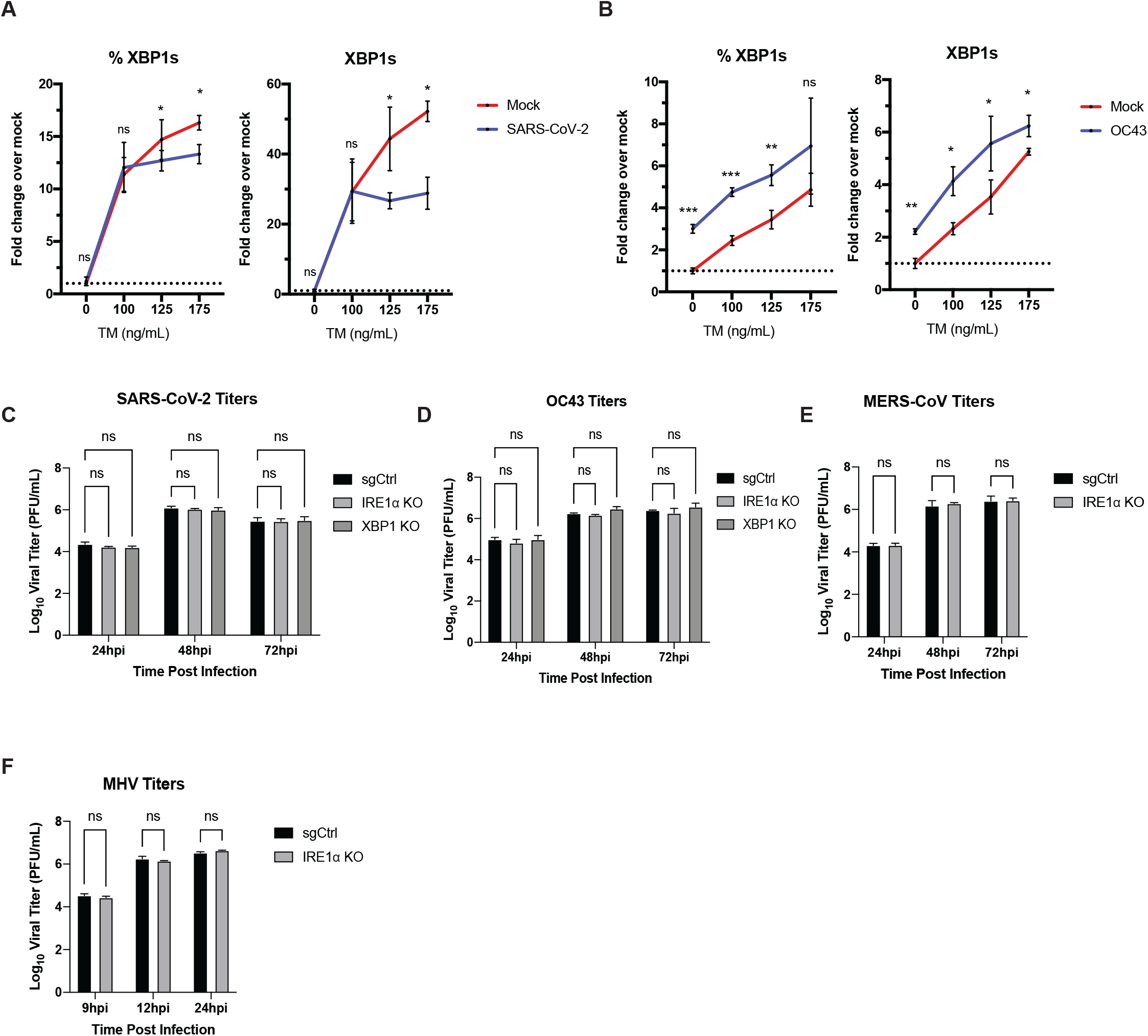
SARS-CoV-2 inhibits IRE1α-mediated XBP1 splicing under ER stress and does not require IRE1α for replication. (A&B) A549-ACE2 cells were mock infected or infected (in triplicate) with SARS-CoV-2 (MOI=3) (A) or OC43 (MOI=1) (B) for 24 hours prior to treatment with low doses of tunicamycin (100-175 ng/mL) for 6 hours. Total RNA was harvested and used to quantify the relative %XBP1s and XBP1s expression by RT-qPCR. C_T_ values were normalized to 18S rRNA and expressed as fold-change over mock displayed as 2^−Δ(ΔCt)^. Technical replicates were averaged, the means for each replicate are displayed as ±SD (error bars). Statistical significance (infected compared to mock) was determined by one-tailed, paired t-tests (* = P < 0.05; ** = P < 0.01; *** = P < 0.001; ns = not significant). (C-F) Infection of CRISPR/Cas9-edited IRE1α KO A549 cells with different coronaviruses. Experiments were performed using sgControl or IRE1α KO or XBP1s KO (where indicated) A549 cells stably expressing viral receptors: A549-ACE2 (OC43 or SARS-CoV-2), A549-DDP4 (MERS-CoV) and A549-MHVR (MHV). Cells were infected (in triplicate) with SARS-CoV-2, MERS-CoV, OC43, or MHV at a MOI of 1. At the indicated times, supernatants were collected and infectious virus quantified by plaque assay. Values are means ± SD (error bars). Statistical significance was determined by two-way ANOVA (* = P < 0.05; ** = P < 0.01; ns = not significant). Data shown are from one representative of at least two independent experiments.

### Betacoronaviruses do not require IRE1α for replication

Given the presumed importance of IRE1α/XBP1s to expand the ER and maintain protein folding during viral replication, and the interesting differences we observed between SARS-CoV-2 and the other betacoronaviruses, we next explored the consequences for its inhibition on the replication of each virus. To determine whether IRE1α activity is required for replication and propagation of MHV, OC43, MERS-CoV or SARS-CoV-2, we utilized CRISPR/Cas9 gene editing to knock out IRE1α in A549 cell lines expressing receptors for each coronavirus (Figure S3 A-F). Surprisingly, we did not observe any significant differences in the capability of all tested coronaviruses to replicate in cells lacking IRE1α (Figure 6C-F). These results suggest IRE1α is neither essential nor inhibitory for coronavirus replication in these cells. Since SARS-CoV-2 does not lead to IRE1α-mediated XBP1 splicing, we also tested replication of SARS-CoV-2 and OC43 does in XBP1s KO cells (Figures 6C&D and S3G). Consistently, there was no detectable effect of XBP1s KO on SARS-CoV-2 or HCoV-OC43 replication in A549-ACE2. Together, these results demonstrate that none of the coronaviruses tested require the activation IRE1α/XBP1 pathway for optimal replication.

### Loss of IRE1α expression causes robust alterations in gene expression, including reduced interferon signaling, following SARS-CoV-2 infection

To gain insight into the role of IRE1α in regulating betacoronaviruses, we conducted RNA sequencing analysis of wildtype or IRE1α knockout A549-ACE2 cells infected with either SARS-CoV-2 or OC43, compared to mock infected cells. Infections of A549-ACE2 cells were carried out at 33C to enable direct comparison of the two viruses [OC43 replication is significantly more robust at 33C compared to 37C, while SARS-CoV-2 replicates to a similar extent at both temperatures (Figure S4A)]. Principal component analysis showed a modest change in cellular gene expression upon OC43 infection of wildtype cells relative to SARS-CoV-2, which caused a robust alteration in gene expression (Figure 7A). In contrast to uninfected or OC43-infected cells, loss of IRE1α significantly impacted host gene expression in SARS-CoV-2-infected A549 cells (Figure 7A,B). Clustering analysis of RNA-seq data revealed 6 distinct clusters altered upon loss of IRE1α related to key cellular functions, including chromatin organization (Cluster 1), mRNA metabolism and processing (Cluster 2) and protein translation (Cluster 3) (Figure 7B: S5A). Detailed analysis of the IRE1α-mediated UPR pathway confirms activation by OC43 infection that is inhibited upon loss of IRE1α (Figure S4C-E). In contrast, minimal change in this pathway was observed in SARS-CoV-2-infected cells, consistent with previous results in this study. Loss of IRE1α also appears to alter other elements of the UPR in SARS-CoV-2-infected cells, including some genes in the PERK and ATF6 pathways (Figure S6), which may reflect compensatory effects on the UPR in an attempt to control proteostasis in the absence of IREα (35-37). Strikingly, we observed significantly lower induction of some interferon stimulated genes (ISGs) during SARS-CoV-2 infection of IRE1α KO cells (Figure 7D, S4F, S5B). We have previously reported that SARS-CoV-2 induces type I and type III IFN signaling and ISGs in multiple cell types (3). Interestingly, OC43 infection did not induce notable IFN or ISG responses with or without IRE1α expression, so we were unable to make the same observations with this virus (Figure 7D). To confirm these results, we performed RT-qPCR on representative IFN and ISG genes that we have previously reported to be upregulated during SARS-CoV-2 infection (3). Consistent with our RNA-seq data, we observed significantly lower induction of ISGs such as OAS2, MX1, and IFIT1 during SARS-CoV-2 infection of cells lacking IRE1α expression at both 37 C (Figure 7E) and 33 C (Figure S4F). These data suggests that IRE1α may play a role in augmenting IFN signaling, while not being necessary for ISG induction, in SARS-CoV-2 infected cells. Our data taken together lead us to propose the model shown in Figure 8.

**Figure 7.**
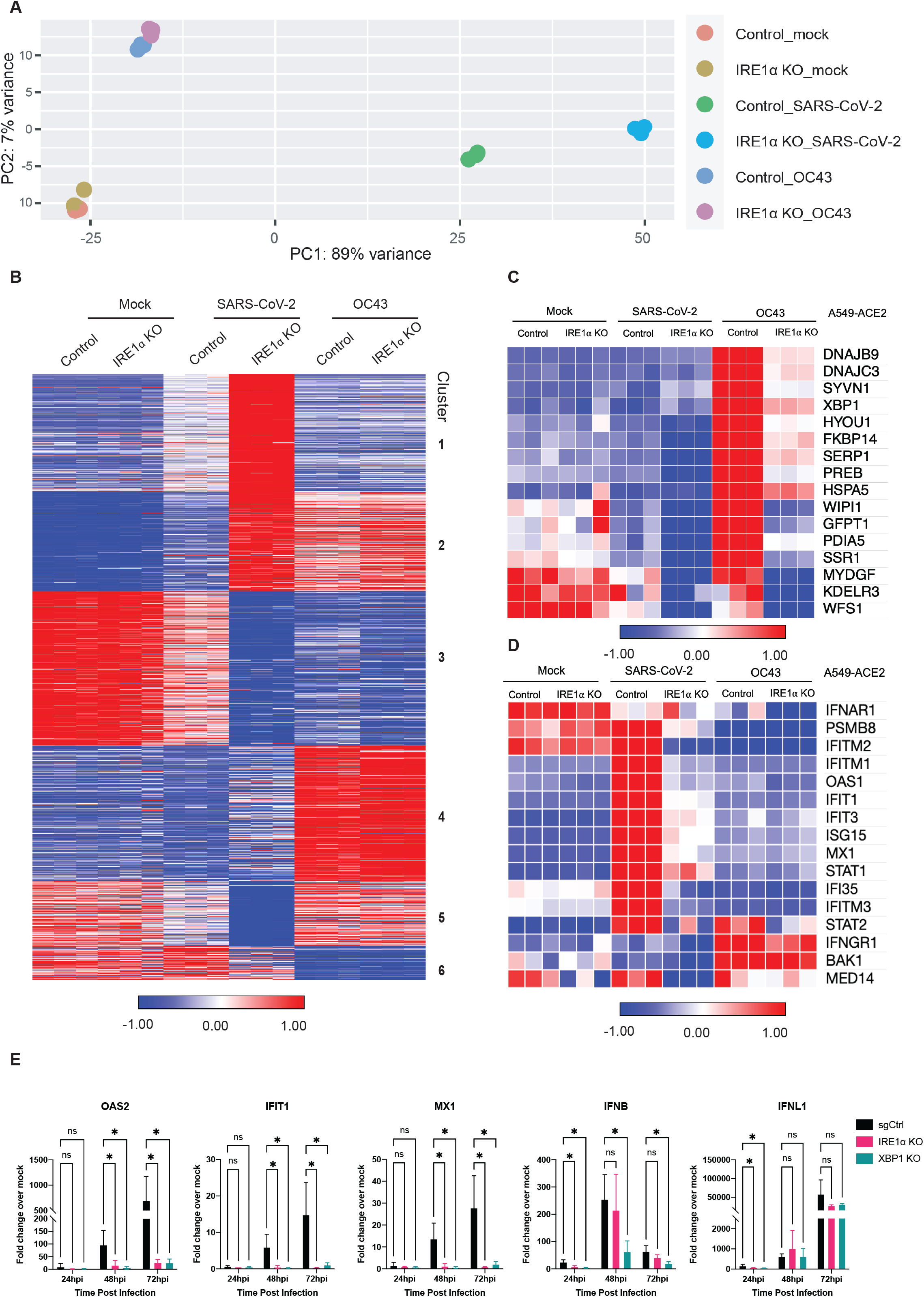
IRE1α promotes the induction of interferon stimulated genes upon SARS-CoV-2 infection. (A-E) A549-ACE2 CRISPR/Cas9-edited IRE1α KO or control cells were mock infected or infected (in triplicate) with SARS-CoV-2 or OC43 (MOI=1) for 48 hours. All infections were performed in the same culture conditions at 33C. Total RNA was harvested and RNA sequencing was performed as described in Materials and Methods. (A) Principal component analysis (PCA) of RNA-seq data from samples in triplicate. The first and second principal components (PC1 and PC2) of each sample are plotted. (B) Heatmap of normalized expression levels of the 5000 most variable genes across all samples were plotted and K-means clustering was used to divided genes into six clusters based on expression patterns among different treatment conditions. (C-D) Heatmap of normalized expression levels from RNA-seq of ER stress IRE1α mediated genes (C) or interferon stimulated genes (D) for all treatment conditions. (E) Total RNA was used to quantify and validate expression of ISGs by RT-qPCR. C_T_ values were normalized to 18S rRNA and expressed as fold-change over mock displayed as 2^−Δ(ΔCt)^. Technical replicates were averaged, the means for each replicate are displayed as ±SD (error bars). Statistical significance (infected compared to mock) was determined by Ordinary one-way ANOVA (* = P < 0.05; ** = P < 0.01; *** = P < 0.001; **** = P < 0.0001 ns = not significant).

**Figure 8.**
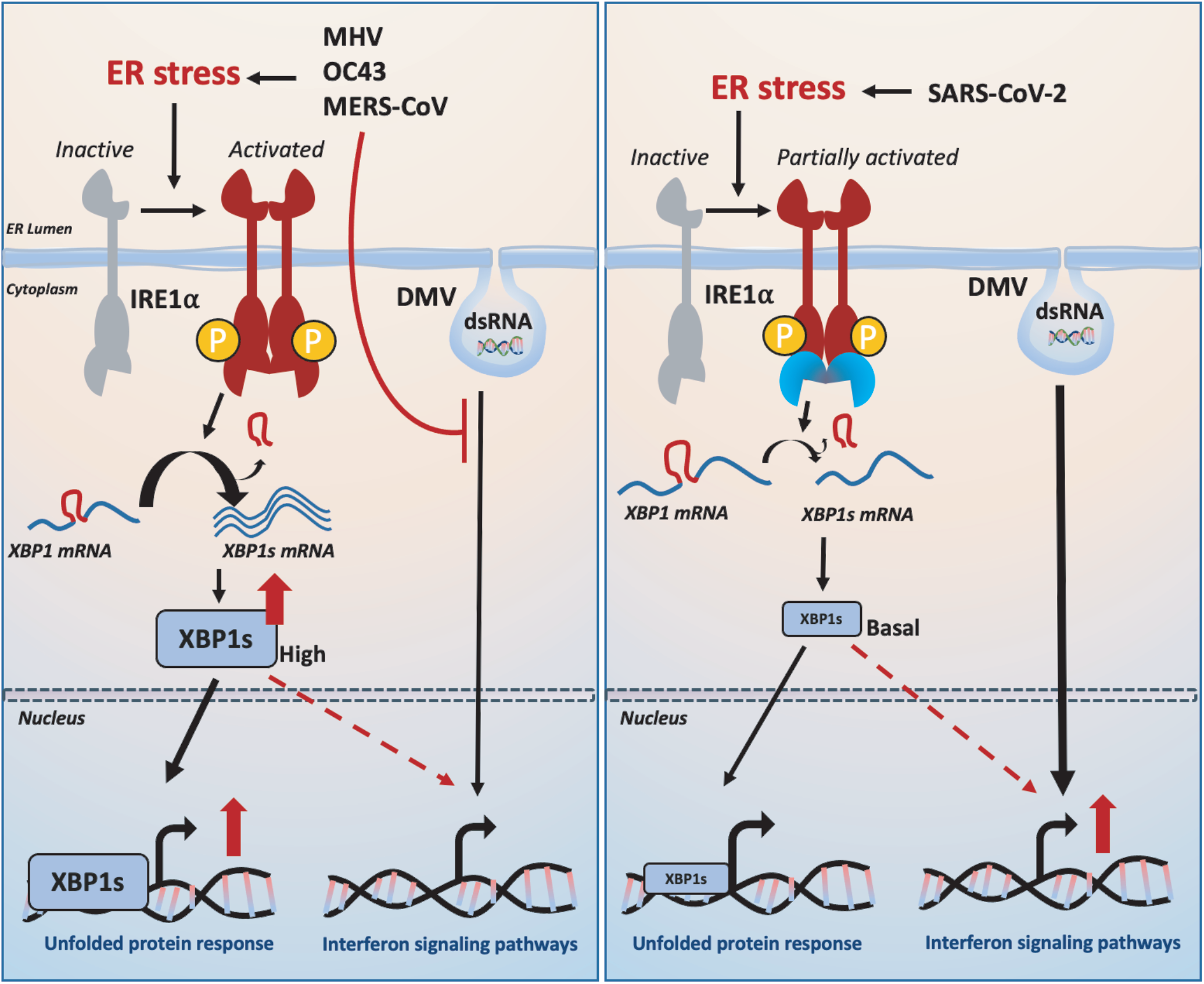
Model of betacoronavirus activation of the IRE1α/XBP1 pathway and downstream effects on interferon signaling. MHV, OC43 and MERS-CoV infection induces ER stress that leads to IRE1α autophosphorylation and downstream IRE1α RNase mediated XBP1 splicing producing XBP1s. In contrast, SARS-CoV-2 infection only partially activates IRE1α through autophosphorylation but prevents the activation of the RNase activity. XBP1s maintains a low basal level upon SARS-CoV-2 infection. MERS, OC43 and MHV efficiently antagonize dsRNA induction of IFN signaling. In contrast, SARS-CoV-2 allows dsRNA induction of some IFN signaling and basal XBP1s potentiates the induction of IFN signaling upon SARS-CoV-2 infection.

## DISCUSSION

Human respiratory betacoronavirusese initiate infection in the upper respiratory tract and have the potential to cause life-threatening pneumonia as a result of infection and inflammation of the lower respiratory tract. The host response to severe infection with CoV is associated with marked dysfunction in the distal lung (alveolar) epithelium, which includes disruption of barrier function, dysregulated immune responses, transcriptomic reprogramming to a transitional cell state, and senescence (38, 39).

To better understand the host epithelial response to CoV, we systematically compared the activation of the IRE1α/XBP1 pathway of the UPR during infection with betacoronaviruses in lung-derived A549 and Calu-3 cells lines and iPSC-derived AT2 cells. We employed three human viruses, each from a different betacoronavirus lineage: OC43 (embeco), SARS-CoV-2 (sarbeco) and MERS-CoV (merbeco), and included the model murine coronavirus MHV, an embecovirus. We found a striking difference between the host response to SARS-CoV-2 and the other three viruses. OC43, MHV and MERS-CoV all activated the canonical IRE1α/XBP1 pathway in both A549 and Calu-3 cell lines as evidenced by phosphorylation of IRE1α (Figure 2), XBP1 mRNA splicing (Figures 3&4) and induction of DNAJB9 (Figure 3), a target of XBP1s. Additionally, MERS-CoV was observed to induce IRE1α/XBP1 activation in iAT2 cells (Figure 5). In contrast, while SARS-CoV-2 also promoted autophosphorylation of IRE1α, there was no evidence of XBP1s, indicating that the pathway was only partially activated and suggesting that the IRE1α kinase was active while the XBP1 splicing RNase activity was not. The differential splicing of XBP1 mRNA during SARS-CoV-2 and MERS-CoV infection was also observed in iPSC-derived AT2 cells, confirming the results in a more physiologically relevant system (Figure 5). The difference among these viruses is surprising as all of them encode highly conserved replicase and structural proteins that promote ER membrane rearrangements and challenge the ER folding capacity, respectively (26). We had originally hypothesized that these conserved genes would induce similar stress on the ER and lead to UPR activation. Instead, our data suggest that that SARS-CoV-2 actively prevents XBP1 splicing (Figure 6A&B). Consistent with this idea, a recombinant SARS-CoV lacking the E protein (rSARS-CoV-ΔE) was reported to induce more XBP1 splicing as well as induction of UPR genes compared to parental wild type virus (40).

To investigate the importance of IRE1α for coronavirus replication, we evaluated replication of each of the betacoronaviruses in IRE1α KO A549 cells compared to parental wild type cells. In contrast to influenza (41), all of the betacoronaviruses examined were able to replicate efficiently in the absence of IRE1α signaling, consistent with a previous report of the gammacoronvirus IBV (25). This raises interesting possibilities for the role of IRE1α during coronavirus infection. As previously stated, IRE1α can produce both cytoprotective (through XBP1s) and destructive responses (via RIDD and JNK/p38 signaling) depending on the extent of the encountered stress. It seems likely that coronavirus infection would induce extensive and prolonged ER stress, which may push IRE1α beyond the initial pro-recovery responses and towards a pro-apoptotic response. Indeed, our data reveal that, at least with MERS-CoV and SARS-CoV-2 infection, IRE1α phosphorylation is readily detectable by 24hpi and remains steady throughout the course of infection (Figure S1A&B). Additionally, unlike what has been observed with chemically induced ER stress (30, 42), IRE1α phosphorylation does not appear to attenuate at any point during coronavirus infection, again suggesting a hyperactive and destructive outcome. As stated above, destruction of cells, in particular AT2 cells in the lung, may contribute to pathogenesis during coronavirus infection. However, SARS-CoV-2 appears to limit the downstream consequences of IRE1α activation, most notably XBP1 splicing via its RNase activity, and thus may be protected from this destructive phenotype. MERS-CoV may induce apoptosis redundantly in the UPR, as it has been reported that MERS-CoV induces and benefits from apoptosis mediated by the PERK arm of the UPR (21, 43).

To further probe the impact of IRE1α signaling on host gene expression following coronavirus infection, we performed RNA sequencing analysis of wildtype or IRE1α knockout A549-ACE2 cells infected with either SARS-CoV-2 or HCoV-OC43. IRE1α deletion significantly reduced the expression of genes downstream of XBP1s during OC43 infection, as expected, with otherwise only modest changes in overall gene expression. In contrast, genetic ablation of IRE1α significantly impacted host gene expression in SARS-CoV-2-infected A549 cells. The two most dramatic effects that appear to be specific to SARS-CoV-2 relate to chromatin organization and protein folding and transport. Effects on mRNA metabolism and processing are also observed for SARS-CoV-2 and, more modestly, for OC43. Finally, protein translation is down-regulated in both OC43 and SARS-CoV-2-infected cells but, in the latter case, occurs primarily upon loss of IRE1α. Taken together, these results suggest that IRE1α plays a key role in mediating changes in host cell gene transcription and protein production caused by SARS-CoV-2.

We found here that deletion of IRE1α modestly blunted the induction of some but not all ISGs by SARS-CoV-2 infection. In contrast, OC43 was not observed to induce significant levels of IFN or ISG mRNAs in either WT or IRE1α KO cells. The mechanism by which loss of IRE1α activity during SARS-CoV-2 infection dampens the induction of interferon signaling remains to be determined. It has been reported that the UPR can precede and prime innate immune signaling in flavivirus-infected cells (44). XBP1s has been found upstream of IFNα and IFNβ transcription and may work through binding upstream cis-acting enhancer elements (45, 46). Moreover, XBP1s can directly bind and transcriptionally activate IL-6, TNFα and other inflammatory cytokines (47). It is possible that a low level of background XBP1 splicing may occur during SARS-CoV-2 infection, which could contribute to these responses. Independent of its RNase activity, the autophosphorylated cytoplasmic domain of IRE1α can oligomerize and serve as a scaffold that recruits TRAF2, JNK, ASK, Nck, and other molecules that can lead to varied signaling outputs (48, 49). Therefore, the ability of SARS-CoV-2 to prevent full IRE1α activation might dampen inflammatory signaling and prevent detection and elimination by the immune system in an intact organism. However, it is important to note that the diminution of ISG expression in the absence of IRE1α is small for most ISGs, and SARS-CoV-2 still induces IFN and IFN signaling to a greater extent than OC43 in IRE1α KO cells. Thus, the significance of IRE1α dependent IFN signaling is not clear and will be a subject of future investigation.

Overall, despite the lack of apparent virus replication defects with IRE1α deficiency, further characterization of the repertoire of betacoronavirus induced IRE1α signaling is warranted, including contributions to cytokine production, apoptosis, and pro-inflammatory responses. While we initially investigated this pathway from the perspective of the impact on virus replication, future studies should examine effects of IRE1α activation on the host, including inflammation and cell death through the JNK and p38 MAPK signaling scaffolded by IRE1α (16) and/or RIDD, as a consequence of prolonged IRE1α activation (11, 50). These responses could be particularly important in AT2 cells, which must rely on the UPR to maintain proteostasis in the face of the challenge from the biosynthesis and secretion of surfactant proteins (51). Dysregulation of these responses by coronavirus infection could promote AT2 cell reprogramming, epithelial apoptosis, alteration of surfactant components in alveoli, and the rampant inflammation associated with severe coronavirus infection (52-54). Finally, the UPR response is complex and made up of the PERK and ATF6 pathways in addition to IRE1α, and signals from all three of these pathways almost certainly integrate into the final outcome of an infected cell.

We recently reported that SARS-CoV-2 and MERS-CoV also diverge in their activation and antagonism of the double-stranded RNA induced host cell innate immune responses, another early innate response to viruses (3). While MERS-CoV actively antagonizes type I and type III interferon production and signaling, the oligoadenylate ribonuclease L (OAS/RNase L) system and the protein kinase R (PKR) pathway, SARS-CoV-2 activates OAS/RNase L, PKR and induces a low level of IFN and ISG expression (3, 4). Here, we observed that OC43 infection did not lead to the induction of IFN or ISGs (Figure 7D), and we have shown previously that OC43 encoded accessory proteins NS2, antagonizes of activation of the OAS/RNase L pathway (55). Activation of these pathways during MERS-CoV mutant infection significantly reduces virus replication (56), while SARS-CoV-2 can tolerate the innate responses activated during infection (3).

Considering the differences we have observed between betacoronaviruses with innate immune responses and now IRE1α activation and signaling, it is striking that MERS-CoV and SARS-CoV-2 are reciprocal in what they activate and antagonize. To optimize replication, coronaviruses must likely strike a balance in the cellular responses they antagonize, tolerate, or benefit from. Supporting this, our data suggest that IRE1α influences ISG induction during infection. It is intriguing to consider if MERS-CoV tolerates this by antagonizing IFN and ISG induction, while SARS-CoV-2 instead limits IRE1α activity. Future studies should examine the synergy between innate immune responses and the UPR during coronavirus infection, and how perturbations on one side may change viral replicative capacity, tropism, and spread. Understanding how signals from each one of these pathways are integrated into viral replication and cell fate decisions during coronavirus infection may illuminate new therapeutic strategies for combating emerging betacoronaviruses.

## MATERIALS AND METHODS

### Cell lines

Human A549 cells (ATCC CCL-185) and its derivatives were cultured in RPMI 1640 (Gibco catalog no. 11875) supplemented with 10% FBS, 100 U/ml of penicillin, and 100 μg/ml streptomycin (Gibco catalog no. 15140). African green monkey kidney Vero cells (E6) (ATCC CRL-1586) and VeroCCL81 cells (ATCC CCL-81) were cultured in Dulbecco’s modified Eagle’s medium (DMEM; Gibco catalog no. 11965), supplemented with 10% fetal bovine serum (FBS), 100 U/ml of penicillin, 100 μg/ml streptomycin, 50 μg/ml gentamicin (Gibco catalog no. 15750), 1mM sodium pyruvate (Gibco catalog no. 11360), and 10mM HEPES (Gibco catalog no. 15630). Human HEK 293T cells (ATCC) were cultured in DMEM supplemented with 10% FBS. Human Calu-3 cells (ATCC HTB-55) were cultured in DMEM supplemented with 20% FBS without antibiotics. Mouse L2 cells(57) were grown in DMEM supplemented with 10% FBS, 100U/mL of penicillin, 100μg/mL streptomycin, 10nM HEPES, 2mM L-glutamine (Gibco catalog no. 25030081), and 2.5μg/mL Amphotericin B (Gibco catalog no. 15290).

A549-DPP4 (4), A549-ACE2 (3) and A549-MHVR (4) cells were generated as described previously. A549-ACE2 cells, used in Figure 3I&J, Figure 4, Figure 6, and Figure S3 were a kind gift of Benjamin TenOever, Mt Sinai Icahn School of Medicine. CRISPR-Cas9 knockout cell lines were generated using lentiviruses. Lentivirus stocks were generated by using lentiCRISPR v2 (Addgene) with single guide RNA (sgRNA) targeting IRE1α sequences: Version1 (V1):CGGTCACTCACCCCGAGGCC, version (V2): TTCAGGAAGCGTCACTGTGC, version (V3): CGGTCACTCACCCCGAGGCC; or XBP1 sequence: TCGAGCCTTCTTTCGATCTC. The infected A549-ACE2 cells were polyclonally selected and maintained by culture in media supplemented with 4 μg/mL puromycin for 1 week.

iPSC- (SPC2 iPSC line, clone SPC2-ST-B2, Boston University) derived alveolar epithelial type 2 cells (iAT2) were grown and infected as previously described (3). In brief, cells were differentiated and maintained as alveolospheres embedded in 3D Matrigel in CK+DCI media, as previously described (58). For generation of 2D alveolar cells for viral infection, alveolospheres were dispersed into single cells, then plated on pre-coated 1/30 Matrigel plates at a cell density of 125,000 cells/cm2 using CK+DCI media with ROCK inhibitor for the first 48h and then the medium was changed to CK+DCI media at day 3 and infected with either mock infected or infected with MERS-CoV or SARS-CoV-2 at a MOI of 5.

### Viruses

SARS-CoV-2 (USA-WA1/2020) was obtained from BEI Resources, NIAID, NIH or provided by Natalia Thornburg, World Reference Center for Emerging Viruses and Arboviruses (Galveston, Texas), and propagated in VeroE6-TMPRSS2 cells. The genome RNA was sequenced and found to be identical to GenBank: MN985325.1. Recombinant MERS-CoV was described previously (1) and propagated in VeroCCL81 cells. SARS-CoV-2 and MERS-CoV infections were performed at the University of Pennsylvania or at the Howard Taylor Ricketts Laboratory (HTRL) at Argonne National Laboratory (Lemont, IL), in biosafety level 3 laboratories under BSL-3 conditions, using appropriate and approved personal protective equipment and protocols. OC43 was obtained from ATCC (VR-1558) grown and titrated on VeroE6 cells at 33C or on A549-mRuby cells as described (59). MHV-A59 (5, 60) was propagated on A549-MHVR cells or on murine 17CL-1 cells.

### Viral growth kinetics and titration

SARS-CoV-2 and MERS-CoV infections and plaque assays were performed as previously described (1, 5). In brief, A549 cells were seeded at 3×10^5^ cells per well in a 12-well plate for infections. Calu-3 cells were seeded similarly onto rat tail collagen type I coated plates (Corning #356500). Cells were washed once with PBS before infecting with virus diluted in serum free media – RPMI for A549 cells or DMEM for Calu-3 cells. Virus was absorbed for 1 hour (A549 cells) or 2 hours (Calu-3 cells) at 37 degrees Celsius before the cells were washed 3 times with PBS and the media replaced with 2% FBS RPMI (A549 cells) or 4% FBS MEM (Calu-3 cells). At the indicated timepoints, 200μL of media was collected to quantify released virus by plaque assay and stored at −80 degrees Celsius. Infections for MHV growth curves were performed similarly in BSL-2 conditions. For OC43 infections, similar infection conditions and media were used, however virus was absorbed, and the infections incubated at 33C rather than 37C.

Plaque assays were performed using VeroE6 cells for SARS-CoV-2 and OC43; VeroCCL81 cells for MERS-CoV; and L2 cells for MHV. SARS-CoV-2 and MERS-CoV plaque assays were performed in 12-well plates at 37C. OC43 and MHV plaque assays were performed in 6-well plates at 33C and 37C, respectively. In all cases, virus was absorbed onto cells for one hour at the indicated temperatures before overlay was added. For SARS-CoV-2, MERS-CoV, and OC43 plaque assays, a liquid overlay was used (DMEM with 2% FBS, 1x sodium pyruvate, and 0.1% agarose). A solid overlay was used for MHV plaque assays (DMEM plus 2% FBS, 1x HEPES, 1x glutamine, 1x Fungizone, and 0.7% agarose). Cell monolayers were fixed with 4% paraformaldehyde and stained with 1% crystal violet after the following incubation times: SARS-CoV-2 and MERS-CoV, 3 days; OC43, 5 days; MHV, 2 days. All plaque assays were performed in biological triplicate and technical duplicate.

### Pharmacologic agents

KIRA8 was purchased at >98% purity from Chemveda Life Sciences India Pvt. Ltd. For use in tissue culture, KIRA8 stock solution was prepared by dissolving in DMSO. Tunicamycin (cat. #T7765) and thapsigargin (cat. #T9033) were purchased at >98% purity from Sigma. For use in tissue culture, tunicamycin and thapsigargin stock solutions were prepared by dissolving in DMSO.

### Immunoblotting

Cells were washed once with ice-cold PBS and lysates harvested at the indicated times post infection with lysis buffer (1% NP-40, 2mM EDTA, 10% glycerol, 150mM NaCl, 50mM Tris HCl, pH 8.0) supplemented with protease inhibitors (Roche complete mini EDTA-free protease inhibitor) and phosphatase inhibitors (Roche PhosStop easy pack). After 5 minutes, lysates were incubated on ice for 20 minutes, centrifuged for 20 minutes at 4°C and supernatants mixed 3:1 with 4x Laemmli sample buffer (Bio-rad 1610747). Samples were heated at 95°C for 5 minutes, then separated on SDS-PAGE, and transferred to PVDF membranes. Blots were blocked with 5% nonfat milk or 5% BSA and probed with antibodies (table below) diluted in the same block buffer. Primary antibodies were incubated overnight at 4°C or for 1 hour at room temperature. All secondary antibody incubation steps were done for 1 hour at room temperature. Blots were visualized using Thermo Scientific SuperSignal chemiluminescent substrates (Cat #: 34095 or 34080).

| Primary Antibody | Antibody species | Blocking buffer | Dilution | Catalog number |
| --- | --- | --- | --- | --- |
| Phospho-IRE1α | rabbit | 5% BSA | 1:1000 | Abcam (EPR5253) |
| IRE1α (14C10) | rabbit | 5% milk/TBST | 1:1000 | Cell Signaling Technology 3294S |
| XBP1 | mouse | 5% milk/TBST | 1:1000 | Biolegend 9D11A43 |
| GAPDH (14C10) | rabbit | 5% milk/TBST | 1:2000 | Cell Signaling Technology 2118S |
| SARS-CoV-2 N | rabbit | 5% milk/TBST | 1:2000 | GTX135357 (Gentex) |
| MERS-CoV N | mouse | 5% milk/TBST | 1:2000 | 40068-MM10 (Sino Biological) |
| OC43 N | rabbit | 5% milk/TBST | 1:2000 | 40643-T62 (Sino Biological) |

### RNA sequencing

A549 cells expressing the MERS-CoV receptor DPP4 (4) were cultured in 10% FBS RPMI media. At 70% cell confluence, cells were washed once with PBS before being mock infected or infected with MERS-CoV (EMC/2012) at MOI = 1. Virus was absorbed for 1 hour at 37 degrees Celsius in serum-free RPMI media. After one hour, virus was removed, cells washed three times with PBS, and 2% FBS RPMI was added. The cells were incubated for another 24 hours or 36 hours, then washed once with PBS and lysed using RLT Plus lysis buffer before genomic DNA removal and total RNA extraction using the Qiagen RNeasy Plus Mini Kit (Qiagen 74134). Three independent biological replicates were performed per experimental condition. RNA sample quality check, library construction, and sequencing were performed by GeneWiz following standard protocols. All samples were sequenced by an Illumina HiSeq sequencer to generate paired-end 150bp reads. Read quality was assessed using FastQC v0.11.2 as described by Andrews, S. (2010) “FastQC: A Quality Control Tool for High Throughput Sequence Data” (http://www.bioinformatics.babraham.ac.uk/projects/fastqc/). Raw sequencing reads from each sample were quality and adapter trimmed using BBDuk 38.73 as described by Bushnell, B at “BBTools software package” (http://sourceforge.net/projects/bbmap). The reads were mapped to the human genome (hg38 with Ensembl V98 annotation) using RNA STAR 2.7.1a(61). The resulting BAM files were counted by featureCounts 1.6.4 to count the number of reads for each gene(62). Differential expression between mock, 24hpi, and 36hpi experimental conditions were analyzed using the raw gene counts files by DESeq2 1.22.1(63). A PCA plot of RNA-seq samples and a normalized gene expression matrix were also generated by DESeq2.

For SARS-CoV-2 and OC43 infections, ACE2-A549 control or IRE1 KO cells were cultured in 10% FBS RPMI to 70% confluence. Cells were washed once with PBS before being mock infected or infected with each virus at MOI = 1 for one hour in serum-free RPMI at 33C. Cells were then washed three times with PBS before 2% FBS RPMI was added. At 48 hours post infection, cells were lysed with RLT Plus lysis buffer before genomic DNA removal and total RNA extraction using the Qiagen RNeasy Plus Mini Kit (Qiagen 74134). Three independent biological replicates were performed per experimental condition. RNA sample quality check, library construction, and sequencing were performed by the University of Chicago Genomics Facility following standard protocols. All samples were sequenced in two runs by a NovaSeq 6000 sequencer to generate paired-end 100bp reads. For each sample, the reads from two flow cells were combined before downstream processing. Quality and adapter trimming were performed on the raw sequencing reads using Trim Galore! 0.6.3 (https://github.com/FelixKrueger/TrimGalore). The reads were mapped to the human genome (UCSC hg19 with GENCODE annotation) and the downstream analyses performed using the same methods as above.

### Host pathway activity analysis of viruses

RNA-seq data from GSE147507(31), GSE168797 (32), GSE144882 (29) and above were used to compare effects of different viruses on host ER stress response. Specifically, Ingenuity Pathway Analysis (IPA) (https://www.qiagenbioinformatics.com/products/ingenuitypathway-analysis) was used to predict activities of related canonical pathways based on host gene expression changes following viral infection. Activation z-scores for every virus and canonical pathway combination were plotted as a heatmap using Morpheus (https://software.broadinstitute.org/morpheus). IPA used the following q-value cutoffs for each dataset to perform the canonical pathway cross comparison: Calu-3 SARS-CoV-2 MOI 2 24hr q < 0.05, NHBE SARS-CoV-2 MOI 2 24hr q < 0.1, A549-ACE2 SARS-CoV-2 MOI 0.2 24hr q < 0.1, A549-ACE2 SARS-CoV-2 MOI 2 24hr q < 0.05, A549-ACE2 SARS-CoV-2 MOI 3 24hr q < 0.01, A549-ACE2 SARS-CoV-2 MOI 1 48hr 33°C q < 0.05, A549-ACE2 OC43 MOI 1 48hr 33°C q < 0.001, A549-DPP4 MERS-CoV MOI 1 24hr q < 0.1, A549-DPP4 MERS-CoV MOI 1 36hr q < 0.01, BMDM MHV-A59 MOI 1 12hr q < 0.1 and over 1-fold up or down-regulated. These cutoffs were implemented due to the limitations set by the IPA software. IPA was also used to overlay gene expression data (log_2_ fold-change) onto the interferon signaling pathway map (Figure S5B).

### Gene expression heatmaps

Expression levels for genes involved in various pathways from RNA-seq data were drawn using Morpheus. For each gene, the normalized expression values of all samples were transformed by subtracting the mean and dividing by the standard deviation. The transformed gene expression values were used to generate the heatmap. For the clustering analysis of RNA-seq experiments for OC43 and SARS-CoV-2-infected A549-ACE2 cells with or without IRE1α, the top 5,000 most variable genes were selected. The normalized gene expression data were analyzed using Morpheus. K-means clustering with 6 clusters was applied to the gene expression data.

### Gene set enrichment analyses

To identify themes across the 6 clusters, functional gene set enrichment analyses for the genes in each cluster were performed using Metascape (64). The following categories were selected for the enrichment analyses: GO Molecular Functions, GO Biological Processes, and KEGG Pathway. Metascape analysis was performed with a minimum *P* value significance threshold of 0.05, a minimum overlap of 10 genes, and a minimum enrichment score of 5. Notable pathways enriched by Metascape from each cluster were summarized in a heatmap using Morpheus. GSEA v4.1.0 (65)was used to perform specific gene set enrichment analyses on Gene Ontology terms : IRE1 mediated unfolded protein response (66, 67); response to type I interferon (68); and response to interferon alpha (69) using the normalized expression data from the RNA-seq experiment for OC43 and SARS-CoV-2-infected A549-ACE2 cells with or without IRE1α.

### Statistical analysis

All statistical analyses and plotting of data were performed using GraphPad Prism software. RT-qPCR data were analyzed by Student’s *t*-test. Plaque assay data were analyzed by two-way ANOVA with multiple comparisons correction. Displayed significance is determined by p-value (P), where * = P < 0.05; ** = P < 0.01; *** = P < 0.001; **** = P < 0.0001; ns = not significant.

### Quantification of XBP1 alternative splicing using RNA-seq data

BAM files produced by RNA STAR were analyzed in Integrative Genomics Viewer 2.9.4 to count the number of XBP1 reads containing the alternative splicing (70). The total number of XBP1 reads were counted by featureCounts. The percentage of XBP1 alternative splicing for each sample was determined by dividing the number of alternatively spliced reads by the number of total XBP1 reads (spliced plus unspliced).

### Quantitative PCR (RT-qPCR)

Cells were lysed with RLT Plus buffer and total RNA was extracted using the RNeasy Plus Mini Kit (Qiagen). RNA was reverse transcribed into cDNA with a High Capacity cDNA Reverse Transcriptase Kit (Applied Biosystems 4387406). cDNA samples were diluted in molecular biology grade water and amplified using specific RT-qPCR primers (see Table below). RT-qPCR experiments were performed on a Roche LightCycler 96 Instrument. SYBR Green Supermix was from Bio-Rad. Host gene expression displayed as fold change over mock-infected samples was generated by first normalizing cycle threshold (C_T_) values to 18S rRNA to generate ΔC_T_ values (ΔC_T_ = C_T_ gene of interest - C_T_ 18S rRNA). Next, Δ (ΔC_T_) values were determined by subtracting the mock infected ΔC_T_ values from the virus infected samples. Technical triplicates were averaged and means displayed using the equation 2^-Δ (ΔCT)^.

Primer sequences list:

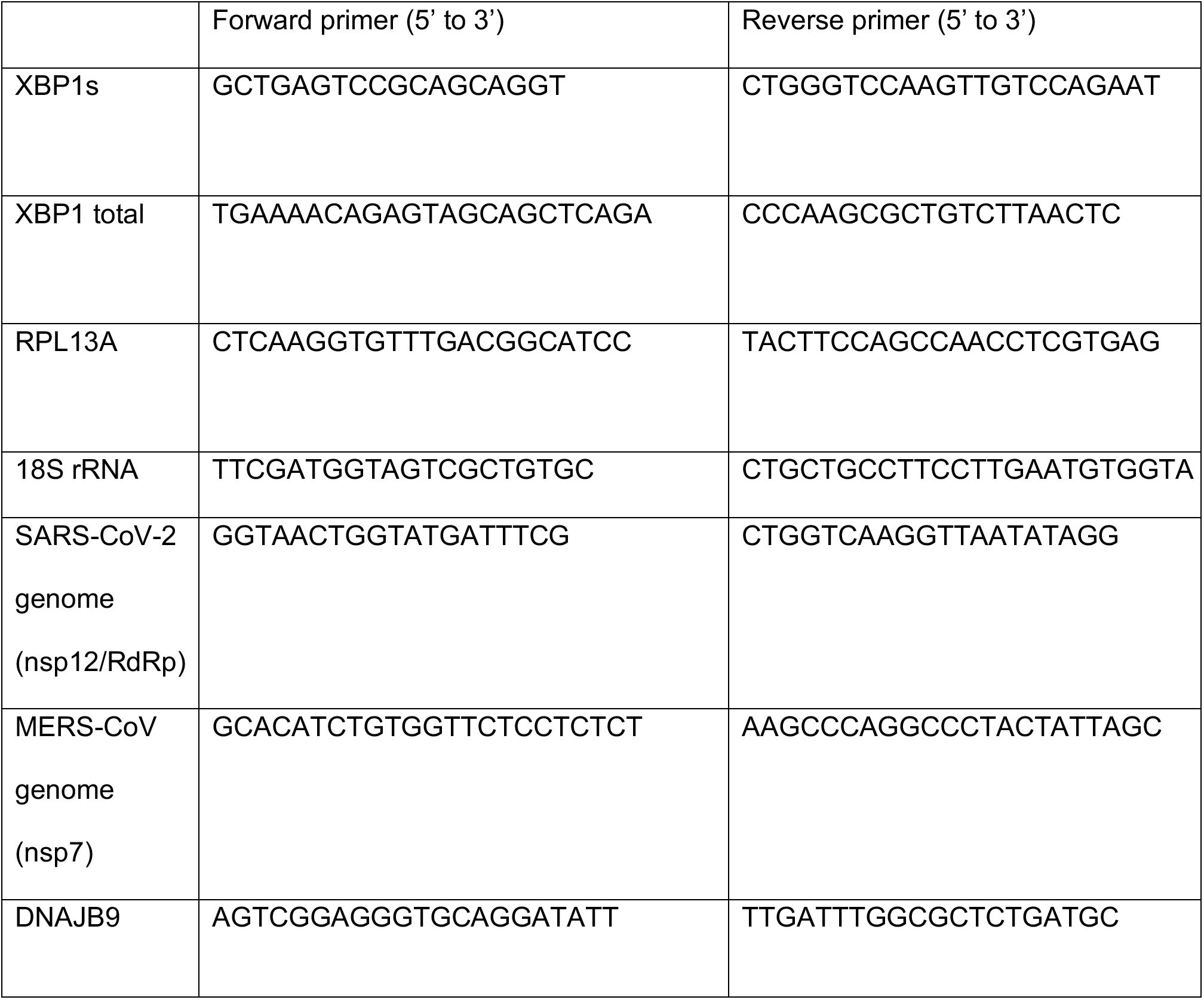

### XBP1 splicing assay by RT-qPCR

RT-qPCR was used to quantify the relative expression of the spliced version of XBP1 (XBP1s) by using specific pairs of primers for human alternatively spliced XBP1 and total XBP1 (primer sequences are described above) as previously described (71). The relative percentage of alternative splicing of XBP1 (%XBP1s) was indicated by calculating the ratio of signals between XBP1s and total XBP1.

### Data Availability

Raw and processed RNA-seq data for MERS-CoV, OC43, and SARS-CoV-2 were deposited into the Gene Expression Omnibus database (GSE193169).

## Acknowledgments

We thank Alejandra Fausto for help with OC43 propagation and titration and Dr. Darrell Kotton (Boston University) and Dr. Rachel Truitt and the Penn iPSC Core for preparation of the iAT2 cells. We thank the members of SARS-CoV-2 host response team in Chicago for stimulating discussions and support: particularly Julian Solway, Rick Morimoto, Nissim Hay, Raymond Roos, and Dominique Missiakas. We thank the University of Chicago Genomics Facility (RRID:SCR_019196) especially Dr. Pieter Faber, for their assistance with RNA sequencing. This work was supported by National Institutes of Health grant R01-AI140442 and supplement for SARS-CoV-2 (S.R.W), R01CA219815 (S.A.O.), R01EY027810 (S.A.O.), U01DK127786 (S.A.O.), R01 GM121735 (M.R.R.); Department of Veterans Affairs Merit Review 2I01BX005411 (M.F.B.&S.R.W.); Penn Center for Research on Coronaviruses and Other Emerging Pathogens (S.R.W); BIG Vision grant from the University of Chicago (M.R.R.); NIH P30 CA014599 (University of Chicago Comprehensive Cancer Center Support grant). DMR was supported in part by National Institutes of Health T32AI055400. DY was supported in part by the Frank W. and Shirley D. Fitch Scholarship Fund.

---

**Supplemental Figure 1.**
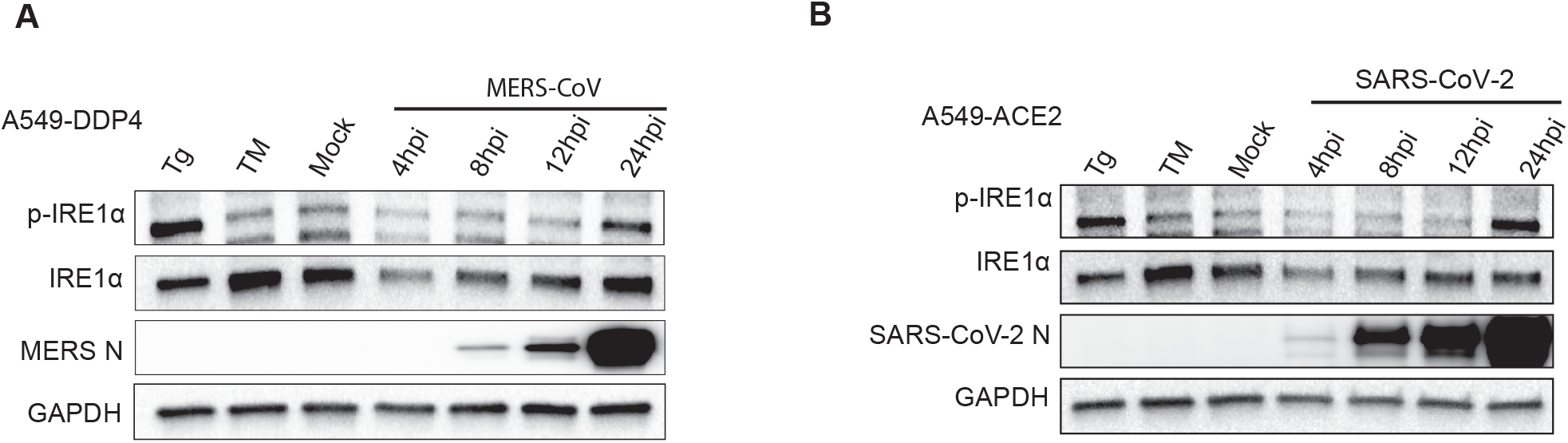
Kinetics of activation of IRE1α phosphorylation during infection with MERS-CoV or SARS-CoV-2. (A-B) A549 cells expressing the indicated viral receptors were mock infected or infected with MERS-CoV (A) or SARS-CoV-2 (B) at a MOI of 5. At the indicated timepoints, total protein was harvested and analyzed by immunoblotting with indicated antibodies. Cells treated with thapsigargin (Tg, 1μM) for 1 hour or tunicamycin (TM, 1μg/ mL) for 8 hours or were used as a positive control for IRE1α phosphorylation and attenuation, respectively. Data shown are from one representative experiment from at least two independent experiments.

**Supplemental Figure 2.**
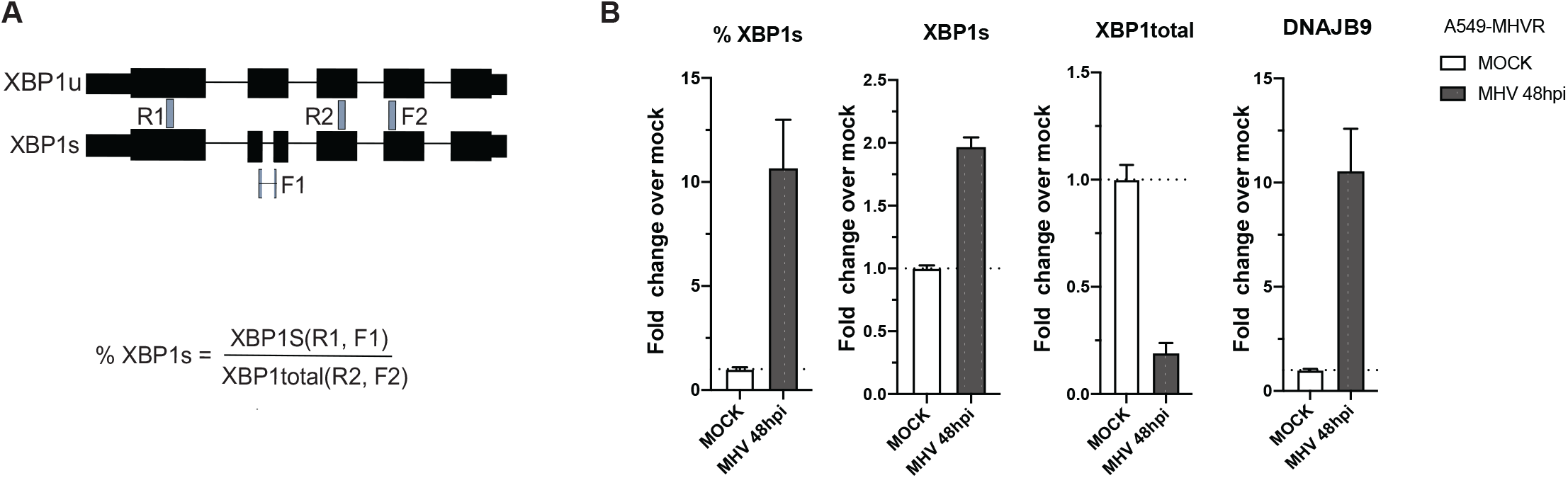
XBP1 is spliced in MHV infected cells. (A) Schematic of method and primer design used to quantify %XBP1. (B) A549-MHVR cells were mock infected or infected with MHV (MOI=0.1). Total RNA was harvested at 48 hours post infection. Relative %XBP1s, XBP1s, total XBP1 and DNAJB9 mRNA expression were quantified by RT-qPCR. C_T_ values were normalized to 18S rRNA and expressed as fold-change over mock displayed as 2^−Δ(ΔCt)^. Technical replicates were averaged, the mean for each biological replicate (n=2) is displayed, ±SD (error bars).

**Supplemental Figure 3.**
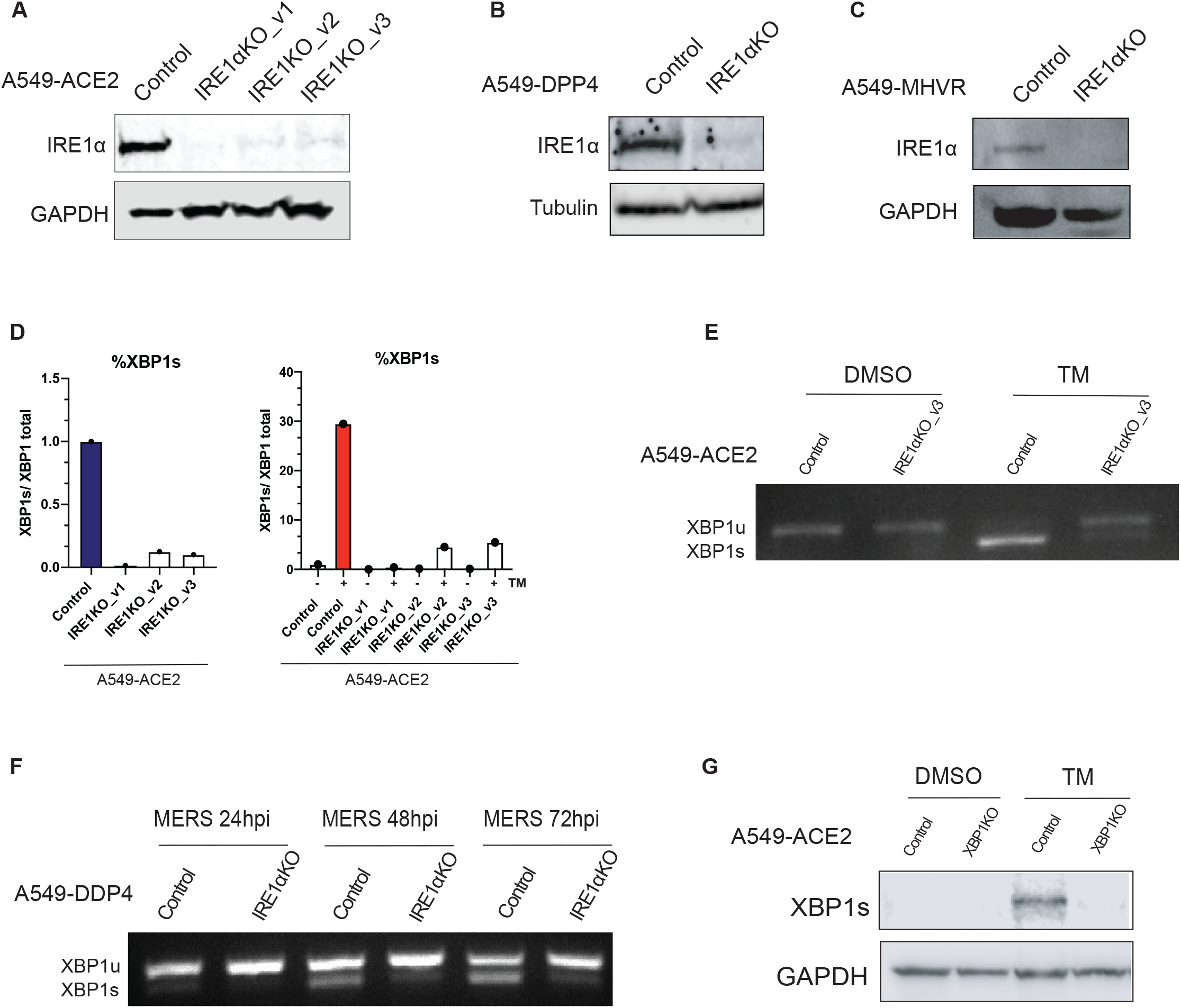
Validation of IRE1α and XBP1 knockout cell lines using CRISPR/Cas9. (A-C) A549 cells expressing the indicated viral receptors subjected to CRISPR/Cas9 editing using different guide RNAs targeting IRE1α were immunoblotted for IRE1α protein to assess knockout efficiency. (D) CRISPR/Cas-9 gene edited IRE1α KO A549-ACE2 cell lines were treated with tunicamycin (500 ng/mL) or DMSO for 6 hours. Total RNA was harvested and %XBP1 quantified by RT-qPCR. Technical replicates were averaged, the means for each replicate displayed. Data shown are one representative experiment from at least three independent experiments. (E) CRISPR/Cas9 gene edited IRE1α KO A549-ACE2 (guide 3) or control A549-ACE2 were treated with tunicamycin (Tm, 1μg/mL) for 8 hours. Total RNA was harvested, reverse transcribed, and amplied for XBP1. XBP1 cDNA product was assayed on an agarose gel to visualize splicing. (F) Control or IRE1α KO A549-DDP4 cells were infected with MERS-CoV (MOI=1). At the indicated time points, total RNA was collected. RT-PCR was performed using primers crossing the XBP1 splicing site. The product was analyzed on an agarose gel to visualize XBP1 splicing. (G) CRISPR/Cas9 gene edited control or XBP1 KO A549-ACE2 were treated with DMSO or tunicamycin (Tm, 1μg/mL) for 6 hours. Lysates were then immunblotted for XBP1s to confirm knockout efficiency.

**Supplemental Figure 4.**
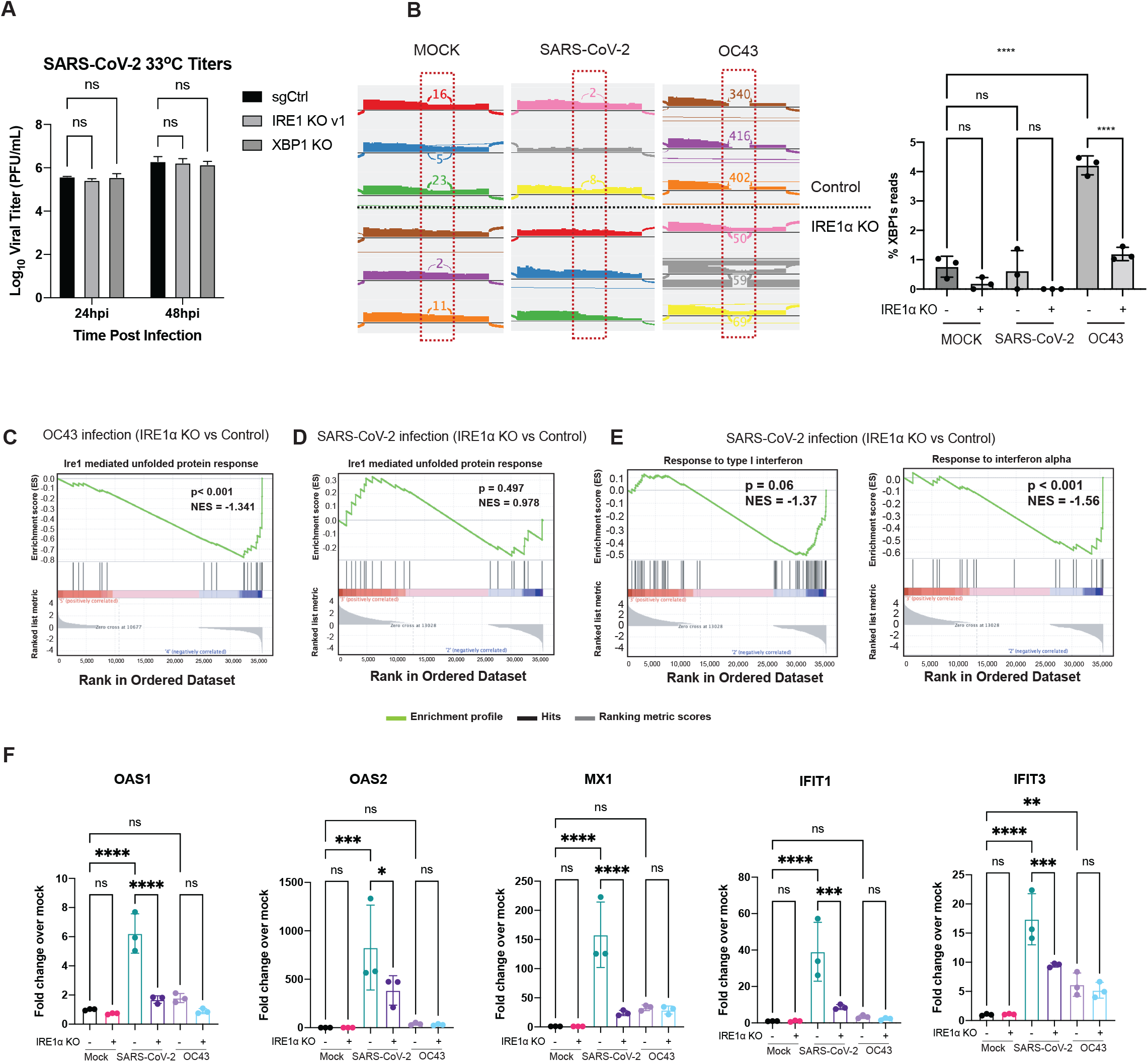
IRE1α promotes the induction of interferon stimulated genes upon SARS-CoV-2 infection. (A) Infection of CRISPR/Cas9-edited IRE1α KO A549-ACE2 cells with OC43 and SARS-CoV-2 (MOI=1) with same culture conditions at 33C. Experiments were performed in triplicate. At the indicated times, supernatants were collected and infectious virus quantified by plaque assay. Values are means ± SD (error bars). Statistical significance was determined by two-way ANOVA (ns = not significant). Data shown are from one representative of at least two independent experiments. (B) Quantification of XBP1 splicing by analyzing RNA-seq data (Figure 7). Reads representing spliced or unspliced XBP1 mRNA were identified based on the presence or absence of the 26-nucleotide intron and quantified. Percentage of XBP1 spliced reads were then plotted. Values are means ± SD (error bars). Statistical significance was determined by ordinary one-way ANOVA. * = P < 0.05; ** = P < 0.01; *** = P < 0.001; **** = P < 0.0001 ns = not significant, adjusted after Tukey’s multiple comparisons test). (C-D) Gene set enrichment analysis (GSEA) of IRE1α mediated unfolded protein response genes with normalized enrichment score (NES) and p-values compared between IRE1α KO and control cells infected with OC43 (C) or SARS-CoV-2 (D). (E) GSEA of genes that belong to GO terms response to type I interferon (left) or response to interferon alpha (right) compared between IRE1α KO and Control cells infected SARS-CoV-2. (F) Infection of IRE1α KO or control A549-ACE2 SARS-CoV-2 (MOI=1) at 33 C. At the indicated times post-infection, total RNA was collected and gene expression quantified by RT-qPCR. C_T_ values were normalized to 18S rRNA and expressed as fold-change over mock displayed as 2^−Δ(ΔCt)^. Technical replicates were averaged, the means for each replicate are displayed as ±SD (error bars). Statistical significance (infected compared to mock) was determined by Ordinary one-way ANOVA (* = P < 0.05).

**Supplemental Figure 5.**
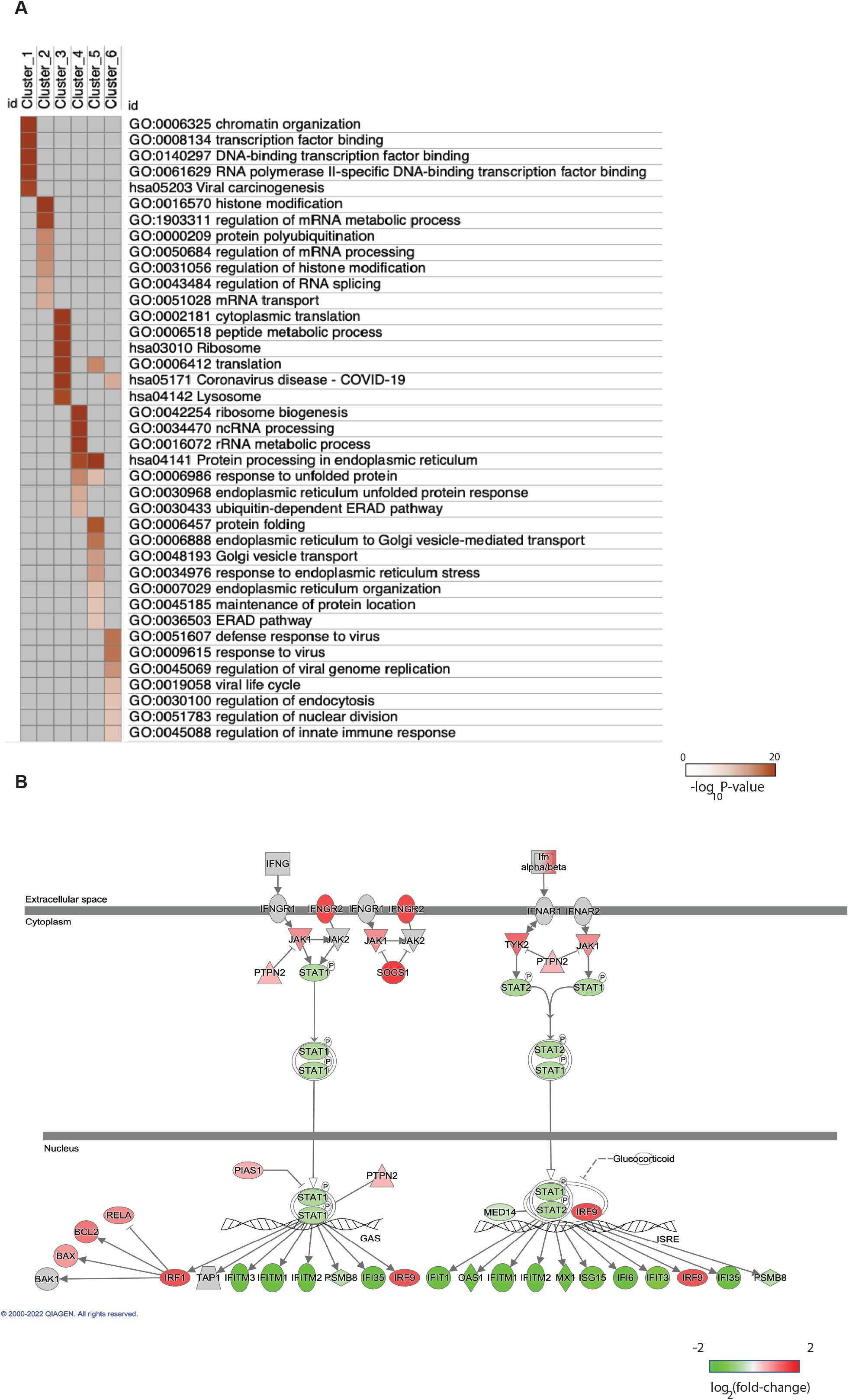
Metascape analysis of SARS-CoV-2 and OC43 infections RNA-seq data. (A) Metascape analyses of genes from six clusters (Figure 7B). GO terms and KEGG pathways (hsa) are shown with -Log10 p-values. (B) Ingenuity-generated interferon signaling pathways analysis compared IRE1α KO over control cells upon SARS-CoV-2 infection from RNA-seq result (Figure 7). Up-regulated genes (red), down-regulated genes (green) or no significant differential expression genes (gray) are shown with color intensity corresponding to log2(fold-change) values from RNA-seq data.

**Supplemental Figure 6.**
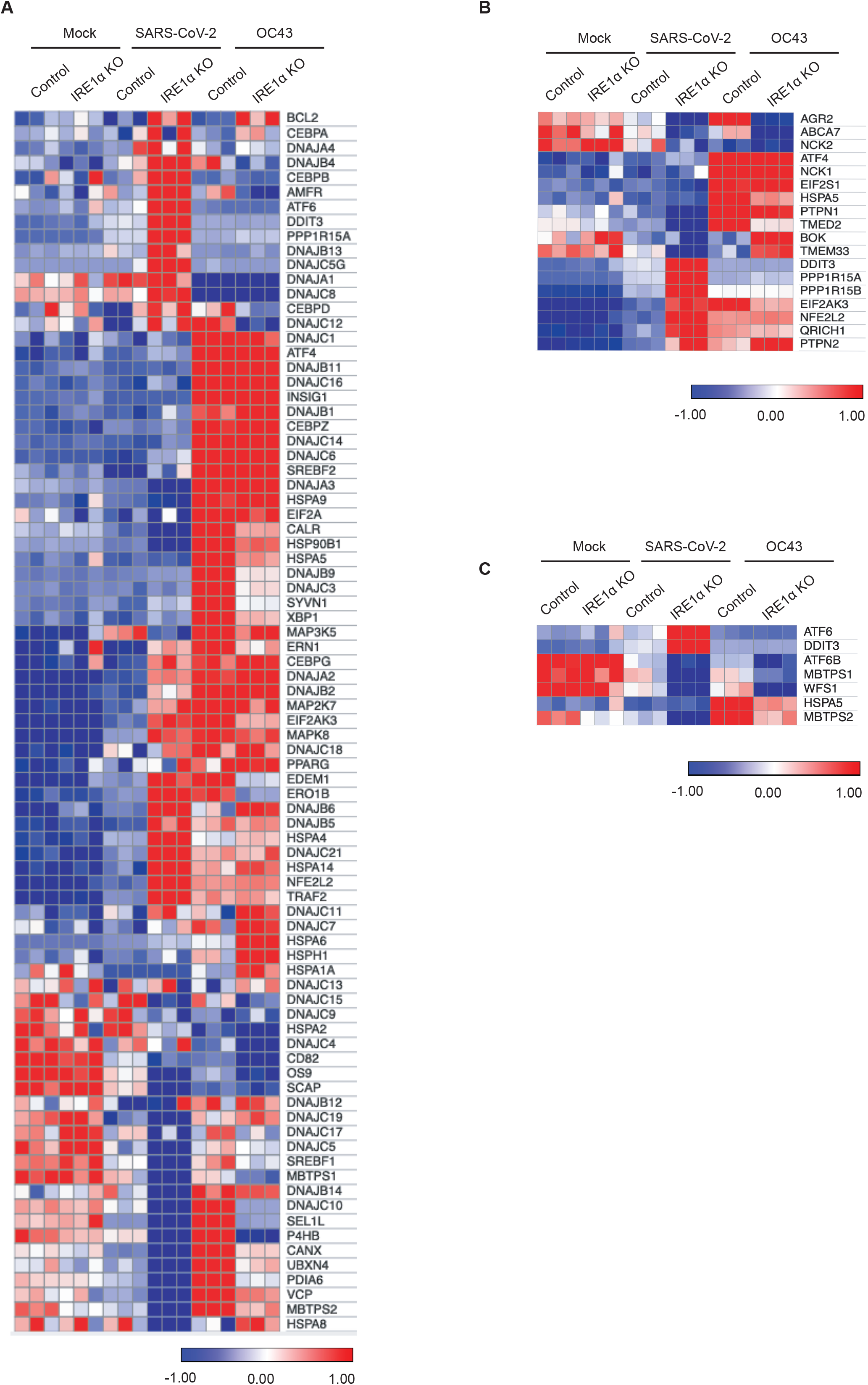
Transcriptomic changes in the host canonical pathway of unfolded protein response upon SARS-CoV-2 and OC43 infection. (A-C) Heatmap of normalized expression levels from RNA-seq (Figure 7) of genes from the canonical pathway of the UPR (A), PERK branch of UPR (B), or ATF6 branch of UPR (C).

